# Weckle is a molecular switch that diverts Toll signalling from innate immunity towards growth by engaging Yki

**DOI:** 10.64898/2026.02.18.706625

**Authors:** Maria Dolores Perez-Sanchez, Guiyi Li, Martin Moncrieffe, Francisca Rojo-Cortés, Karina Malinovska, Emily Sample, Myles Maddick, Marta Moreira, Elizabeth Connolly, Anna Parsons, Roberto Feuda, Nick J. Gay, Alicia Hidalgo

## Abstract

The mechanisms underlying how the brain can switch from a plastic to a degenerative state are unknown. Tolls and Toll-Like receptors (TLRs) could be involved since they underlie both neuroinflammation and structural brain plasticity. Here, we show that the Toll signalling adaptor Weckle (Wek) - a ZAD-Zinc-finger transcription factor - can switch the response from promoting innate immunity or cell death, to inducing growth instead. Myristoylated Wek binds Tolls at the membrane, and concerted PI3K and Toll signalling drive its nuclear translocation. Wek interacts with Yorkie (Yki) – the key regulator of growth restrained by Hippo - enabling its nuclear shuttling. Remarkably, Wek blocks innate immunity. Instead, Wek and Yki together drive glial cell proliferation and growth. Through this mechanism, Toll signalling could promote structural brain plasticity via Wek and Yki downstream. As Tolls and Yki are widespread, these findings could have important implications for understanding also regeneration and cancer.

## INTRODUCTION

The brain changes throughout life, with brain plasticity enabling adaptation and processes such as neuroinflammation driving degeneration. Brain plasticity involves gliogenesis and myelination, neurogenesis, neurite growth, synaptogenesis and circuit remodelling, i.e. a regenerative signature (Birey et al., 2017; Feldman and Brecht, 2005; Gage, 2019; Holtmaat and Svoboda, 2009). Exercise, learning and anti-depressants induce structural brain plasticity, which enhances learning, resilience to stress and wellbeing (Gage, 2019; Troubat et al., 2021; Zhao et al., 2020). By contrast, neuroinflammation leading to neurite, synapse and cell loss, is emerging as a driver of psychiatric conditions (e.g. depression, anxiety) and neurodegenerative diseases (e.g. Alzheimer’s disease) (Bock, 2021; Shi and Yong, 2025; Troubat et al., 2021). What enables the brain to switch between plastic and inflammatory, or regenerative and degenerative, states is unknown. Working this out is essential to understand how the brain works, and how to direct brain and mental health.

Toll and Toll-Like Receptors (TLRs) could be key. They are best known for their universal function in promoting innate immunity across the animals (Gay and Gangloff, 2007; Hoffmann, 2003; Imler and Hoffmann, 2001; Leulier and Lemaitre, 2008). In humans TLRs also underlie neuroinflammation, stroke, and neurodegenerative diseases, and links of TLRs to schizophrenia-like behaviour have been found in mice (Kumar, 2019; Okun et al., 2009; Park et al., 2015; Pascual et al., 2021; Squillace and Salvemini, 2022). In microglia, TLRs drive clearance of pathogens and cell debris resulting from damage, infections and defective proteostasis, but in inflammatory states they also lead to abnormal neurite and synapse pruning and cell loss (Kumar, 2019; Squillace and Salvemini, 2022). In *Drosophila*, activation of Toll-1 by fungi can also drive immune evasion leading to cell loss in the brain (Singh et al., 2025). Fungal infections have also been linked to neuroinflammation and neurodegeneration also in humans (Alonso et al., 2014; Pisa et al., 2015; Seelig et al., 2019). In response to pathogens, Tolls and TLRs activate signalling through the universally conserved adaptor Myeloid differentiation primary response 88 (MyD88) and the downstream pathway kinase Pelle/Interleukin Receptor-Associated Kinase 1 (Irak1), in *Drosophila* and mammals respectively. This results in the phosphorylation of the NFκB inhibitor Cactus/Ikb, thereby promoting its degradation (Gay and Gangloff, 2007; Hoffmann and Reichhart, 2002; Leulier and Lemaitre, 2008). This enables the nuclear translocation of Dif/NFκB, leading to the activation of anti-microbial peptide gene expression to eliminate pathogens, and the up-regulation of pro-inflammatory cytokine gene expression (Gay and Gangloff, 2007; Hoffmann and Reichhart, 2002; Kumar, 2019; Leulier and Lemaitre, 2008; Squillace and Salvemini, 2022). However, both in *Drosophila* and mammals, Tolls and TLRs can also function in the brain independently of immunity (Anthoney et al., 2018). In mammals, TLRs can promote neuronal death or survival; regulate neurogenesis and gliogenesis; modulate pain sensitivity via ion channels; regulate dopaminergic signalling; and are involved in learning, memory and feeding behaviour (Abarca-Merlin et al., 2024; Anthoney et al., 2018; Chen et al., 2019; Donnelly et al., 2020; Li et al., 2021; Okun et al., 2010; Rolls et al., 2007; Sanchez-Petidier et al., 2022; Shechter et al., 2008). However, the functions of mammalian TLRs in the healthy, intact brain remain relatively unexplored. In *Drosophila*, Tolls are receptors for neurotrophin family ligands encoded by the *spätzle (spz)* paralogue gene group. Neurotrophins are the main plasticity growth factors both in the mammalian (including human) and *Drosophila* brains (Lu et al., 2005; McIlroy et al., 2013; Park and Poo, 2013; Sun et al., 2024; Wang et al., 2022; Zhu et al., 2008). At least seven of the nine Toll paralogues are expressed in the adult fruit-fly brain, in overlapping yet complementary domains that demark distinct brain modules and neural circuits (Li et al., 2020). In the absence of infection, Tolls can regulate neuronal survival or death, autophagy, engulfment of dead neurons by glia, adult neurogenesis, synaptogenesis, neural circuit remodelling, dopaminergic signalling, locomotion and long-term memory (Li et al., 2020; Li and Hidalgo, 2021; McIlroy et al., 2013; McLaughlin et al., 2016; McLaughlin et al., 2019; Sakakibara et al., 2023; Singh et al., 2025; Sun et al., 2024; Ulian-Benitez et al., 2017; Zhang et al., 2024). Remarkably, over-expression of Toll-2 induces adult neurogenesis from quiescent progenitor cells that co-express MyD88 and the pan-glial marker Repo, and this adult-induced cell proliferation requires both *weckle (wek)* and *yorkie (yki)* (Li et al., 2020).

Wek is a Toll signalling adaptor (Chen et al., 2006), that has received little attention. It was proposed that Wek can bring MyD88 closer to Toll-1, facilitating its function in dorso-ventral patterning, but not in immunity (Chen et al., 2006). Later in development, Wek enables the adaptor Sterile Alpha and TIR Motif (Sarm) to function downstream of Tolls (Foldi et al., 2017). Sarm is localised at the plasma membrane and in the mitochondria, serving as the highly evolutionarily conserved and only inhibitor of MyD88 and TRIF (Belinda et al., 2008; Carty and Bowie, 2019; Carty et al., 2006; Peng et al., 2010). Sarm can inhibit innate immunity; facilitate JNK signalling to induce cell death; and with its catalytic NADase function, promote NAD breakdown, neurite destruction and Wallerian degeneration (Essuman et al., 2017; Foldi et al., 2017; Mukherjee et al., 2015; Osterloh et al., 2012; Singh et al., 2025). Toll signalling via Wek can promote cell death in multiple contexts by engaging Sarm (Foldi et al., 2017; Singh et al., 2025). In humans, some studies have linked Sarm to nervous system diseases (Gilley et al., 2021; Lin and Hsueh, 2014; Xiang et al., 2022). Despite these important functions, Wek is a zinc-finger protein (ZFP), predicted to also function in the nucleus.

Wek is a ZFP containing a ZAD-domain (Chung et al., 2007). ZFPs are transcription factors that regulate gene expression in the nucleus. There are 15 orthologous groups containing ZAD domains in invertebrates, and 100 ZAD-containing genes in *Drosophila*, including 4 *wek* paralogues (Bonchuk et al., 2021; Chung et al., 2002). However, what genes Wek and its paralogues may regulate as transcription factors is not known. The ZAD domain belongs to a broader family of SCAN, KRAB, ICAROS and BTB domains present in ZFPs throughout animals. These domains evolved for genome defence to control the mobilisation of transposable elements (Ecco et al., 2017). The co-evolution of these ZFPs and the transposons they tamed resulted in them regulating host cell state, including stemness and cell fate (Krystel and Ayyanathan, 2013).

Here, we asked what enables Wek to switch Toll signalling from promoting innate immunity (via MyD88) and promoting cell death (via Sarm) to inducing cell proliferation instead. We show that Wek binds Tolls via their TIR domain, it is located at the plasma membrane by myristoylation and its nuclear translocation depends on phosphorylation by the Toll signalling kinase Pelle and the insulin-signalling effector PI3K. Wek facilitates the nuclear shuttling of Yki. Wek inhibits innate immunity and induces a morphogenetic or regenerative programme instead. We show that in vivo, Wek functions together with Yki to promote glial cell proliferation and growth. This novel mechanism could enable Wek to promote structural brain plasticity and regeneration downstream of Toll signalling.

## RESULTS

### Wek is a zinc-finger protein that directly binds Toll receptors

To model the structure of Wek, we used Uniprot sequence Q9VJN5 as query in AlphaFold. The structure of ZAD proteins is exemplified by the crystallisation of *Drosophila* Grauzone, a ZAD-ZFP involved in germline development (Chung et al., 2007; Jauch et al., 2003). Wek contains a ZAD domain, a disordered region and six zinc finger repeats (Figure 1A,B,C). The Wek-ZAD domain can dimerise (Figure 1D). This was verified by analytical ultracentrifugation of the purified Wek-ZAD domain, which revealed Wek ZAD monomers, dimers and tetramers (Figure 1E). This shows that Wek can dimerize and form larger complexes.

**Figure 1.**
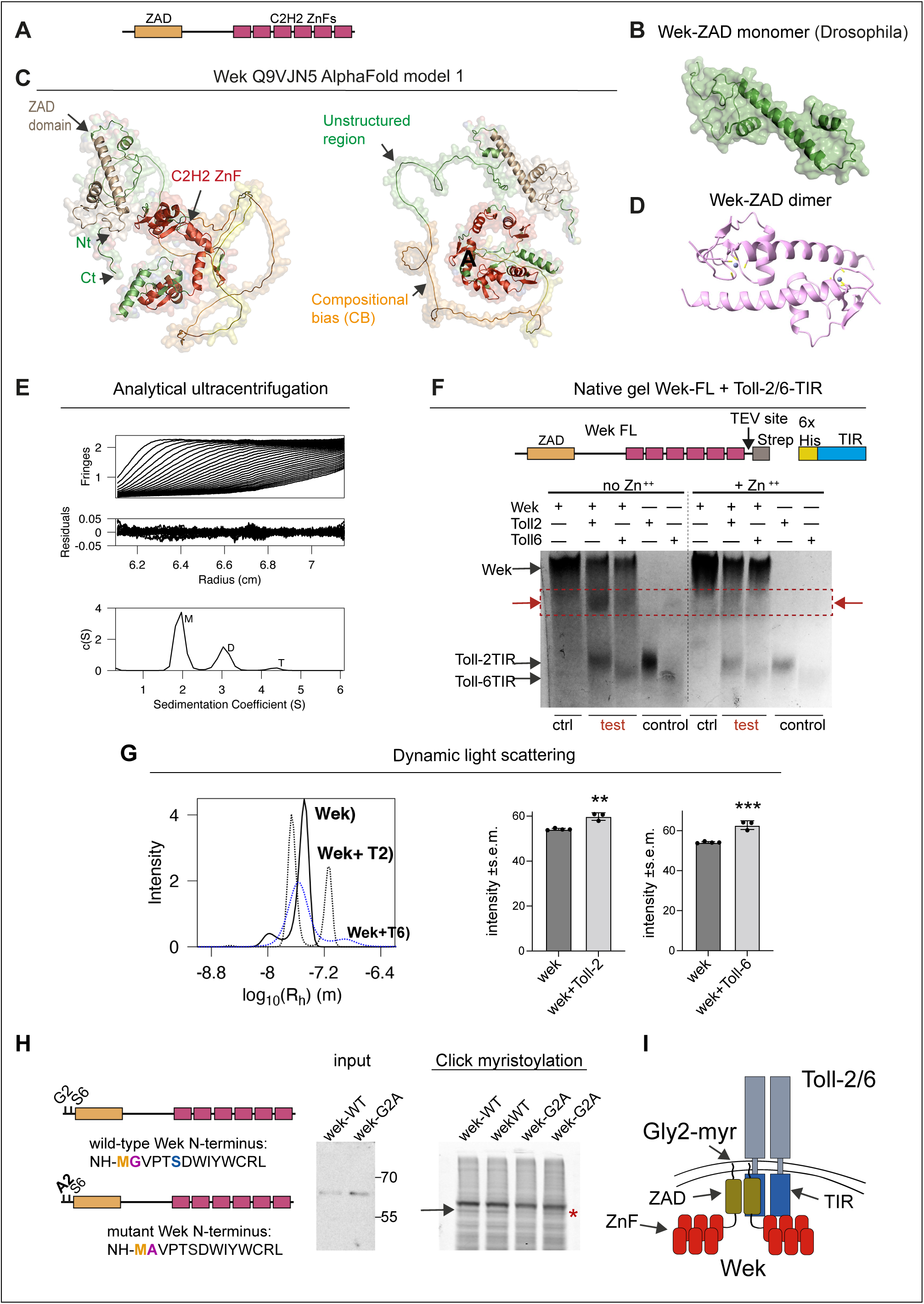
Wek is a ZAD-ZFP that binds Tolls upon myristoylation. **(A)** Diagram of Wek structure**. (B, C, D)** AlphaFold models of protein ID Q9VJN5: **(B)** Wek ZAD domain, **(C)** full-length Wek and **(D)** Wek dimer. Nt: N-terminus and C-t: C-terminus of Wek. C2H2 ZnF: zinc-fingers. **(E)** Analytical centrifugation evidence that Wek-ZAD can form dimers and tetramers. **(F)** Native gel showing that the migration profile of full-length Wek is altered by the addition of the TIR domains of either Toll-2 or Toll-6 (dotted box and arrows). **(G)** Dynamic light scattering evidence that Wek interacts with Toll-2 and Toll-6 (see Supplementary Figure S1). Unpaired Student t tests **p=0.0011, ***p=0.0005, n=3-4. **(H)** Site-directed mutagenesis of *wek* was used to change Gly2 to Ala. Click-myristoylation reaction showing that the Wek band is lost in the *wek-G2A* mutant lanes (arrow indicates wil-type Wek and asterisk the missing band in the mutant). **(I)** Illustration of the association of Wek with the TIR domain of Tolls at the plasma membrane.

Wek is thought to be located at the plasma membrane, and it can bind Tolls, MyD88 and Sarm in co-immunoprecipitation experiments (Chen et al., 2006; Foldi et al., 2017). However, Toll receptors generally interact with other proteins via their TIR domains, which Wek lacks. To further test whether Wek can directly bind Tolls, we expressed and purified full-length (FL) Wek and the TIR domains from both Toll-2 and Toll-6 and ran them together in a native gel. We found a mobility shift when comparing the migration of Wek-FL, Toll-2-TIR and Toll-6-TIR alone vs. when migrating together (Figure 1F). There was no improvement in the presence of Zn++ (Figure 1F). These data show that Wek can bind the TIR domain of Toll-2 and -6 in vitro. To verify this finding, we carried out Dynamic Light Scattering (DLS), whereby particles scatter light differentially based on their size. We found that the DLS profile of purified Wek-FL was modified when it was combined in vitro with both purified Toll-2-TIR or Toll-6-TIR (Figure 1G). Together, these data showed that Wek can directly bind the TIR domain of Toll-2 and -6 at the plasma membrane.

It was intriguing that a zinc-finger transcription factor could be located at the plasma membrane. Cytoplasmic proteins lacking a transmembrane domain can be tethered to the plasma membrane by myristoylation. Wek has the consensus myristoylation sequence at the N terminus, with a Glycine in position 2 followed by a Serine in position 6 (Figure 1H) (Grand, 1989; Towler et al., 1988). We used site directed mutagenesis to mutate the evolutionarily conserved Gly2 into Alanine *(wek-G2A)* and test whether Wek is normally myrystoylated. Wild-type and mutant FLAG-tagged *wek* were expressed in S2 cells and verified by western blot (Figure 1H). Using a click fluorescence myristoylation assay, we found a band in wild-type Wek that was lost in the *wek-G2A* mutant samples (Figure 1H), demonstrating that Wek is myristoylated at Gly2. Altogether, these data show that Wek is myristoylated and thereby tethered to the plasma membrane, where it interacts with Tolls (Figure 1I).

### The subcellular localisation of Wek is regulated by phosphorylation

Wek was reported to be located both at the plasma membrane and in nuclei (Chen et al., 2006). Nuclear shuttling is generally regulated by phosphorylation. Myristoylated proteins can be dissociated from the plasma membrane by phosphorylation at the N-terminus, close to the consensus sequence, possibly Ser6 (Rowe et al., 2006; Thelen et al., 1991). A high-throughput proteome analysis of *Drosophila* had identified phosphorylation of Wek at Ser168 (Zhai et al., 2008) (Figure 2A). To test whether Wek is phosphorylated by Toll signalling, we transfected S2 cells with either *wek-FLAG* alone or *wek-FLAG* plus *Toll-6-HA*, purified Wek and analysed it by mass spectrometry. We had previously shown that over-expression of *Toll-6* in S2 cells is sufficient to induce signalling and that S2 cells naturally express all Toll receptors and their downstream signalling effectors (Foldi et al., 2017). Upon co-expression of *wek-FLAG* and *Toll-6HA,* Wek was phosphorylated at Ser164 (Figure 2B). To confirm this, we co-transfected S2 cells with *wek-FLAG* and the Toll signalling kinase *pelle-V5* and stimulated them with purified DNT-2, the ligand of Toll-6. This also resulted in the phosphorylation of Wek at Ser164 (Figure 2B). These data showed that Toll signalling via Pelle caused the phosphorylation of Wek at Ser164.

**Figure 2.**
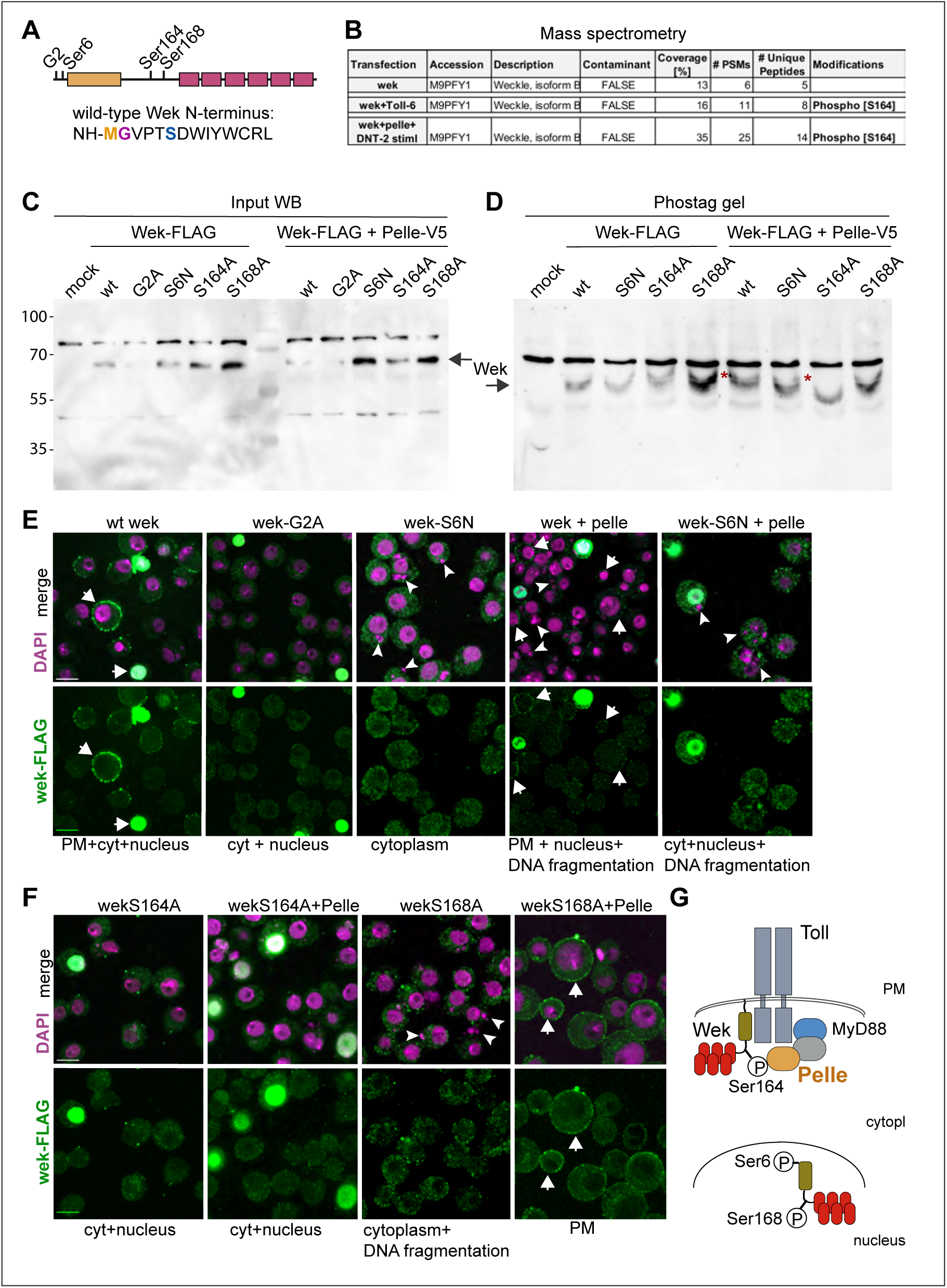
Multiple phosphorylation events regulate the subcellular distribution of Wek. **(A)** Diagram of Glycine2 and Serines in Wek tested. **(B)** Mass spectrometry of purified Wek-FLAG: S2 cells were co-transfected with *wek-FLAG* and *Toll-6HA,* or *wek-FLAG* and *pelle-HA* and stimulated with purified DNT-2, Wek was purified, and found to be phosphorylated at Ser164. This Ser was not phosphorylated in controls. **(C,D)** Phostag gel of *wek* mutant forms expressed in S2 cells, showing: **(C)** western blot input; **(D)** Phostag gel, showing the distinct migration profile of the Wek mutant forms. Asterisks indicate bands modified by the co-transfection with Pelle. **(E,F)** Microscopy from transfected S2 cells showing that site directed mutagenesis of *wek* in *G2A, S164A* and *S168A,* alter the subcellular distribution of Wek, and this is further influenced by *pelle*. Wek visualised with anti-FLAG antibodies (green) and nuclei with DAPI (magenta). **(G)** Illustration showing that phosphorylation at Ser6 and Ser168 are required for the nuclear translocation of Wek. Scale bar: **(E,F)** 10μm

To test whether the subcellular localisation of Wek could be regulated by phosphorylation, we used site-directed mutagenesis to prevent Wek phosphorylation at Ser6 (Serine to Asparagine, S6N), Ser164 (Serine to Alanine, S164A) and Ser168 (Serine to Alanine, S168A) (Figure 2A). After transfecting S2 cells with FLAG-tagged *wek*, we ran a phostag gel that allows the separation of protein phospho-forms. This showed that wild-type Wek is naturally phosphorylated, as multiple bands could be detected, and importantly, the migration profiles of mutant Wek forms differed to that of the controls (Figure 2C,D). Co-transfecting S2 cells with wild-type *wek-FLAG* and *pelle-V5* resulted in an additional band (Figure 2D), consistent with the mass spectrometry data that Pelle phosphorylates Wek. However, there was a reduction in phosphorylation when co-expressing *wek-S6N-FLAG* and *wek-S164A-FLAG* mutant forms, and *pelle-V5* (Figure 2D). These data suggest that Wek phosphorylation at Ser6 by an unknown kinase and at Ser-164 by Pelle facilitated further phosphorylation events. Perhaps phosphorylation at Ser6 brings Wek closer to Pelle, enabling phosphorylation at Ser164, which in turn facilitates phosphorylation of Wek by other kinases.

To test whether phosphorylation influenced the subcellular distribution of Wek, we visualised Wek-FLAG in transfected S2 cells with anti-FLAG antibodies. Wild-type Wek-FLAG was found at the plasma membrane, cytoplasm and nucleus (Figure 2E). By contrast, in *wek^G2A^-FLAG* transfected cells, membrane signal was lost (Figure 2E), confirming that Wek is myristoylated at Gly2. Wek^S6N^-FLAG was predominantly found in the cytoplasm (Figure 2E). Importantly, co-transfection with both wild-type *wek-FLAG* and *pelle-V5* enriched Wek at the membrane and caused nuclear fragmentation (Figure 2E). Nuclear fragmentation is a feature of apoptosis, suggesting that the phosphorylation of Wek by Pelle enabled its apoptotic functions.

Importantly, Wek^S164A^-FLAg - which cannot be phosphorylated by Pelle - was found in the nucleus and cytoplasm, but no longer at the plasma membrane (Figure 2F). This profile was not altered when cells were co-transfected with both *wek^S164A^-FLAG* and *pelle-V5* (Figure 2F). There was no nuclear fragmentation in *wek^S164A^-FLAG* transfected cells, both with and without *pelle-V5* (Figure 2F). Together, these data show that Pelle phosphorylates Wek at Ser164 anchoring it at the plasma membrane, and this facilitates apoptosis.

This is consistent with the fact that Toll-dependent apoptosis requires Wek binding the cell executioner Sarm at the plasma membrane (Foldi et al., 2017; Singh et al., 2025). When we transfected S2 cells with *wek^S168A^,* Wek was excluded from nuclei (Figure 2F). Most remarkably, co-transfection with *pelle-V5* retained *Wek^S168A^-FLAG* at the plasma membrane (Figure 2F). These data suggested that phosphorylations at Ser6 and Ser168 are required for the nuclear translocation of Wek.

Altogether, these data showed that Wek is phosphorylated by Pelle, which retains it at the plasma membrane where it can drive apoptosis. This also enables phosphorylation of Wek by other kinases, driving its nuclear translocation (Figure 2G).

### PI3K signalling regulates the nuclear shuttling of Wek

To find out what factors might drive Wek’s subcellular shuttling, we carried out a pull-down assay followed by mass spectrometry, from S2 cells co-transfected with *wek-FLAG* and *Toll-6HA,* and precipitation of Wek-FLAG (Figure 3A-E, Table S1). This identified factors involved in cell growth (e.g. translation factors, ribosomal proteins, RNA processes, protein folding and processing factors), cell death, cell proliferation, vesicle trafficking, synaptic proteins, and regulation of Wingless, insulin and Hippo signalling. Important hits included (Figure 3E): (1) Sarm, driver of apoptosis and axon destruction, previously known to interact with Wek (Foldi et al., 2017; Singh et al., 2025), serving as a positive control; (2) JNK, which induces apoptosis downstream of Toll, Wek and Sarm (Foldi et al., 2017); (3) 14-3-3e and z, which capture in the cytoplasm proteins that normally shuttle in and out of the nucleus, most particularly Yki (Oh and Irvine, 2008); (4) Yki and Msn, of the Hippo signalling pathway; (5) cell cycle factors, like Cdc5 and Polo; (6) Akt-1, S6K kinase and IGF-II mRNA binding protein (Imp) of the insulin signalling pathway.

**Figure 3.**
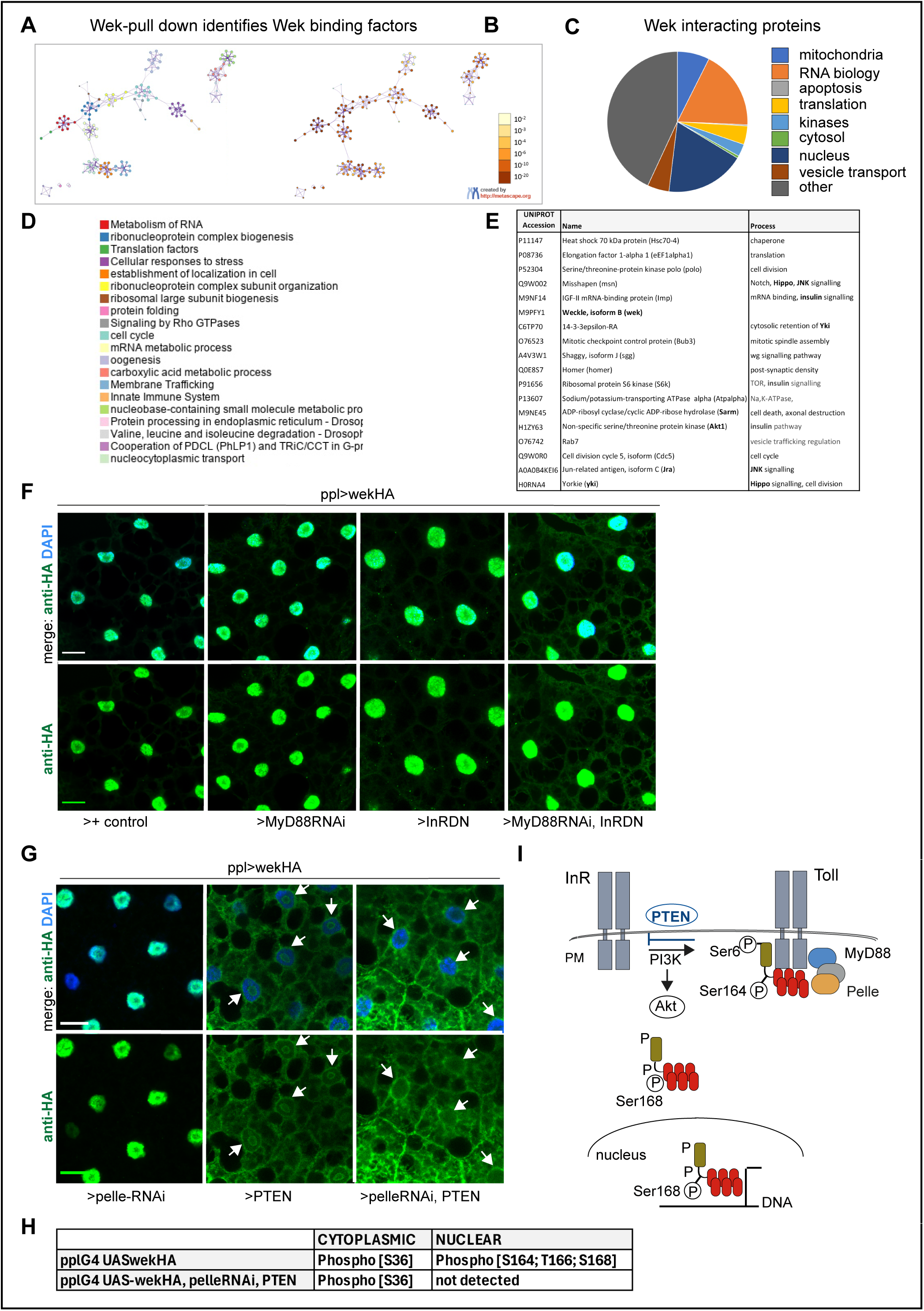
Combined insulin/PI3K and Toll signalling drive the intracellular localisation of Wek. **(A, B)** Wek-FLAG pull-down hits, from transfected S2 cells, shown as Metascape diagram; **(B)** p values; **(D)** colour key. See Table S1. **(C)** Wek interacting proteins. **(E)** Subset of Wek interacting proteins. **(F)** Over-expressed *wek-HA* in the fat body with *pplGAL4* is distributed exclusively to nuclei, visualised with DAPI. Co-expression with *MyD88RNAiGD9716* and *InR^DN-K1409A^*got Wek-HA partially out of nuclei, as it was also present in cytoplasm and plasma membrane. **(G)** Co-expression of *wek-HA* together with *PTEN* increased the cytoplasmic retention of Wek, although Wek remained in nucleoli. Co-expression with *PTEN* and *pelle-RNAi^TRiP.HMS04458^*prevented the nuclear distribution of Wek, and it was enriched in cytoplasm and plasma membrane instead. Arrows indicate nuclei, visualised in blue with DAPI. **(H)** Cell fractionation and mass spectrometry evidence that wild-type Wek phosphorylated at Ser164 and Ser168 localises to nuclei, and that concerted phosphorylation by Pelle and PI3K is required for the nuclear shuttling of Wek, as these phosphorylations are lost with *pelle-RNAi, PTEN* over-expression *(ppl>wek-HA, pelle-RNAi^TRiP.HMS04458^, PTEN).* **(I)** Diagram illustrating how insulin and Toll signalling together regulate the nuclear shuttling of Wek**. (F,G)** Scale bars: 20μm. See Table S11 for genotype details and sample sizes.

The fat body naturally has both active Toll and insulin signalling, and over-expressed Wek is distributed in fat body nuclei (Chen et al., 2006). Furthermore, fat body cells are large and ideal to visualise the sub-cellular distribution of proteins. Thus, we asked whether concerted regulation of Wek by insulin and Toll signalling may drive its nuclear translocation. To this aim, we tested if knock-down of Toll and insulin signalling would prevent the nuclear import of Wek. We over-expressed tagged *wek-HA* in the fat body (with *pplGAL4*) and it was only present in nuclei (Figure 3F). Over-expression of *wek-HA* together with RNAi knock-down of the Toll signalling adaptor MyD88 *(ppl>wek-HA, MyD88RNAi^GD9716^)* did not affect the distribution of Wek (Figure 3F). Over-expression of *wek-HA* together with a *dominant negative form of the insulin receptor (ppl>wek-HA, InR^DN^)* resulted in the slight presence of Wek-HA in the cytoplasm (Figure 3F). However, when we over-expressed *wek-HA* together with *InR^DN^* as well as *MyD88RNAi^GD9716^*, Wek was found in nuclei, cytoplasm and plasma membrane (Figure 3F). To further verify this, we over-expressed *wek-HA* whilst at the same time knocking-down *pelle* to prevent Toll signalling and over-expressing *PTEN* to inhibit PI3K downstream of InR signalling *(ppl>wek-HA, pelle-RNAi^TRiP^ ^HMS04458^, PTEN).* Knock-down of *pelle* with *pelle-RNAi^TRiP^ ^HMS04458^* did not affect the nuclear distribution of Wek (Figure 3G). Over-expression of *PTEN* alone was sufficient to drive Wek out of the nuclei, although nucleolar signal remained (Figure 3G). In contrast, co-expression of *pelle-RNAi^TRiPHMS04458^* and *PTEN* caused a dramatic and complete exclusion of Wek-HA from the nuclei (including the nucleoli), and a pronounced increase in its distribution in the cytoplasm and at the plasma membrane (Figure 3G). Using cell fractionation followed by mass spectrometry, we could identify Wek phosphorylation at Ser164 and Ser168 in the nuclear, but not cytoplasmic fraction (Figure 3H). Phosphorylation at these sites were lost upon *pelle RNAi^TRiP^ ^HMS04458^* knock-down and *PTEN* over-expression *(ppl>wek-HA, pelle-RNAi^TRiP^ ^HMS04458^, PTEN*) (Figure 3H). This shows that PI3Kinase signalling, presumably via Akt downstream, phosphorylates Wek at Ser168 driving its nuclear translocation.

To conclude, these data showed that concerted phosphorylation by the Toll signalling kinase Pelle and a kinase downstream of PI3K, is required for the nuclear shuttling of Wek (Figure 3I).

### Wek and Pelle facilitate the nuclear translocation of Yki

Wek pull-down also brought down Yki as a potential interacting protein (Figure 3). This was interesting, as Toll-2-dependent neurogenesis requires both *wek* and *yki* (Li et al., 2020). Yki is a co-transcriptional activator negatively regulated by the Hippo and AMPK (in the CNS) signalling pathways, which phosphorylate Yki to inhibit its nuclear translocation (Davis and Tapon, 2019; Gailite et al., 2015; Huang et al., 2005; Zheng and Pan, 2019). Yki can be located at the plasma membrane bound by Merlin and Expanded, or retained in the cytoplasm by 14-3-3, and when released, it can translocate into the nucleus (Davis and Tapon, 2019; Huang et al., 2005; Zheng and Pan, 2019). In the nucleus, Yki promotes cell proliferation, inhibits apoptosis, promotes cell growth and activates neural stem cells (Davis and Tapon, 2019; Fan et al., 2023; Parker and Struhl, 2015; Poon et al., 2016; Zheng and Pan, 2019). Yki does not directly bind DNA but instead it is a co-activator of transcription factors, such as Scalloped (Sd) in *Drosophila* (Davis and Tapon, 2019; Zheng and Pan, 2019). Yki pull-down with Wek (Figure 3) suggested the intriguing possibility that Yki may also function together with Wek. To explore this, we first validated the pull-down with co-immunoprecipitation from S2 cells (Figure 4A). This showed that precipitating Wek-HA with magnetic beads brought down Yki, which we visualised with anti-Yki antibodies (Figure 4A). To test a reciprocal relationship and whether it could also occur in vivo, we over-expressed both *yki-V5* and *pelle-HA* in the fat body (with *tubGAL80ts, pplGAL4),* precipitated Yki-V5, and this brought down Wek from the nuclear cell fraction (Figure 4B). Importantly, using mass spectrometry, we found that nuclear Wek was phosphorylated at Ser164 and Ser168 (Figure 4B), consistent with our previous findings (Figures 2 and 3). This confirmed that Wek and Yki can physically interact.

**Figure 4.**
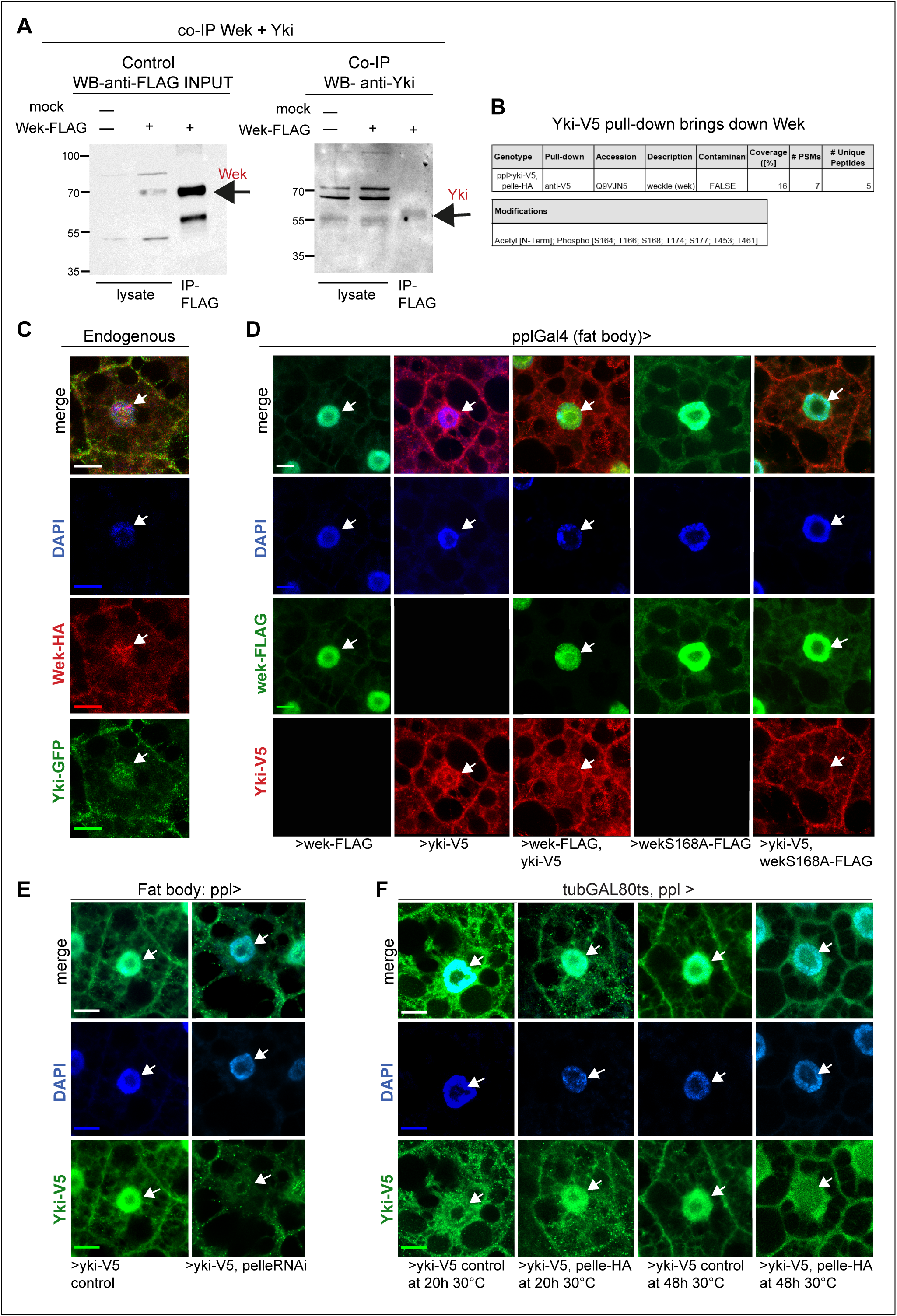
Wek interacts with Yki and facilitates its nuclear translocation. **(A)** Co-immuno-precipitation of Wek and Yki. S2 cells transfected with *wek-FLAG* and precipitated with anti-FLAG brought down Yki, visualised with anti-Yki antibodies. **(B)** Yki-V5 pull-down from larvae *(ppl>yki-V5)* larvae brought down Wek from the cellular nuclear fraction, where Wek was phosphorylated at Ser164 and Ser168. **(C)** Endogenously tagged Wek-HA and Yki-GFP and visualised with anti-HA and anti-GFP antibodies, reveal co-localisation in fat body nuclei visualised with DAPI (genotype: *w; wek-HA/+; yki-GFP/+).* **(D)** Excess Wek-HA and Yki-V5 in the fat body from co-over-expression *(ppl>wek-HA, yki-V5)* reduced the nuclear import of Yki. Mutant *wek^S168-^FLAG* was present in cytoplasm and plasma membrane. Over-expression of mutant *wek^S168A^-FLAG* prevented the nuclear import of Yki *(ppl> wek^S168A^-FLAG, yki-V5)*. **(E)** *pelle-RNAi^TRiP.HMS04458^*knock-down prevented the nuclear translocation of Yki *(ppl>yki-V5, pelle-RNAi^TRiP.HMS04458^).* Penetrance: 28% n= 54 nuclei with no Yki-V5 vs control: 0% n=50 nuclei, Fisher’s Exact test p<0.0001. **(F)** Over-expression of *pelle-HA (tubGAL80^ts^, ppl>yki-V5, pelle-HA)* did not interfere with the nuclear import of Yki (20h at 30°C, penetrance: 82% n= 51 nuclei with high Yki within vs control: 71% n=45 nuclei, Fisher’s Exact test n.s. p=0.2284), but over time it caused Yki’s exclusion from nuclei (48h at 30°C, penetrance: 100% n= 75 nuclei with low Yki vs control: 25% n=40 nuclei, Fisher’s Exact test p<0.0001). **(C,D,E,F)** Arrows indicate nuclei. Scale bars: 15μm. See Table S11 for genotype details and sample sizes.

Next, we asked what consequence the interaction between Wek and Yki might have in their subcellular distribution, looking in fat body cells due to their large size. To first test if Wek and Yki naturally colocalise in fat body cells, we generated a *wek-HA* knock-in allele and using anti-HA found Wek to co-localise with endogenously tagged Yki-GFP in fat body cell membranes, cytoplasms and - predominantly - in nuclei (Figure 4C). Next, we asked whether excess of Wek and Yki could affect this distribution by over-expressing tagged *wek-FLAG* and *yki-V5* in the fat body (ie *ppl>wek-FLAG, yki-V5*). Over-expressed Wek-FLAG alone was found predominantly in the nuclei with weak signal in cytoplasm and at the plasma membrane (Figure 4D). Over-expressed Yki-V5 was abundant at the plasma membrane, cytoplasm and nuclei (Figure 4D). By contrast, in cells co-over-expressing both, there was a decrease in the nuclear distribution of Yki-V5 (Figure 4D). As endogenously expressed Wek and Yki co-existed in nuclei, these data suggest that the physical interaction between Wek and Yki impaired their subcellular trafficking, most particularly the nuclear shuttling of Yki. To further test this, we over-expressed mutant *wek^S168^FLAG*, which favours the membrane localisation of Wek in S2 cells (Figure 2F). Over-expression of *wek^S168^FLAG* in vivo *(ppl>wek^S168^FLAG)* did not prevent the nuclear distribution of Wek in fat body cells, but it increased its cytoplasmic and plasma membrane distribution compared to wild-type Wek (Figure 4D). Remarkably, over-expressed Wek^S168^-FLAG prevented the nuclear translocation of Yki-V5 *(ppl>wek^S168^-FLAG, yki-V5),* as no Yki-V5 was found in nuclei nor in nucleoli (Figure 4D). Together, these data showed that Wek facilitates the nuclear translocation of Yki.

We had shown above that the nuclear translocation of Wek requires both Toll and insulin signalling (Figures 2 and 3). Insulin can also drive the nuclear translocation of Yki in neural stem cells (Strassburger et al., 2012). Thus, we wondered whether Toll signalling might influence the subcellular distribution of Yki too. First, as we did for Wek, we asked how over-expression of *PTEN* and *pelle-RNAi* knock-down in the fat body would affect the subcellular distribution of Yki *(ppl>yki-V5, PTEN, pelle-RNAi)*. However, this genotype caused lethality. Instead, we knocked-down *pelle-RNAi* alone (*ppl> yki-V5, pelle-RNAi),* and this could cause a dramatic exclusion of Yki from 28% nuclei (Figure 4E). This suggested that Pelle is required, although not essential, for the nuclear shuttling of Yki. To further test whether Pelle could promote the nuclear translocation of Yki, we over-expressed both *pelle-HA* and *yki-V5* in the fat body *(ppl>pelle-HA, yki-V5),* but this caused lethality. Thus, we used conditional over-expression with *tubGAL80ts*, switching on *pelle* expression for only 20h or 48h at 30°C during larval development *(tubGAL80ts, ppl>ykiV5, pelle-HA*). Over-expression of *pelle-HA* for 20h at 30°C did not compromise the nuclear import of Yki (Figure 4F). However, after 48h at 30°C, nuclei over-expressing *pelle* had reduced Yki levels within (Figure 4F). This was consistent with findings showing that in the context of infection, Pelle promotes Hippo signalling by phosphorylating Cka, leading to nuclear exclusion of Yki (Liu et al., 2016). Together, these data indicate that *pelle* is required for the initial nuclear translocation of Yki, but over time it prevents it.

To conclude, these data show that Wek is a partner of Yki; that Wek facilitates the nuclear translocation of Yki; and that the Toll signalling kinase Pelle can influence the nuclear translocation of both Wek and Yki.

### *wek* switches off innate immunity and induces a regenerative signature

Wek is a zinc-finger protein, a type of transcription factor. Thus, we asked what genes might be regulated by nuclear Wek. We had previously shown that Toll-2 can induce adult neurogenesis via *wek* and *yki* (Li et al., 2020). Interestingly, Toll-2 is associated to PI3K (Tamada et al., 2021), which could facilitate the nuclear functions of Wek. Thus, we sought to identify Wek targets in the brain. As we had shown that over-expressing both Wek and Yki together impaired their sub-cellular trafficking, we over-expressed *wek* alone. To identify genes regulated by Wek in the brain, we over-expressed *wek* in adult flies in all cells expressing the canonical Toll signalling adaptor MyD88 and carried out a bulk RNAseq analysis from heads. We used *tubGAL80ts* to restrict *wek* expression to adult flies only, isolated the heads, and compared their transcriptome to those of control flies *(tubGAL80ts Myd88>wek vs tubGAL80ts Myd88GAL4/+)* (Figure 5A). In *wek* over-expressing heads, there was a remarkable downregulation in the expression of genes involved in innate immunity, mostly of the Toll pathway, including those encoding: pattern-recognition receptors (e.g. *PGRP-SA, GNBP-like 3, GNBP2*), peptidases (e.g. *serpin, SPE, necrotic*) and anti-microbial peptides against Gram+ bacteria and fungi (e.g. *drosomycin, bomanins, metchnikowin*); but also against Gram- (e.g. *attacins*) and viruses (e.g. *nazo*) (Figure 5A, Supplementary Figure S2, Table S2). Up-regulated genes included those involved the cell cycle; gene transposition and chromatin; neuroblast fate; cell growth or axon targeting; and neuronal genes (Figure 5B, Table S3). Most importantly, these data showed that the over-expression of *wek* switched off innate immunity.

**Figure 5.**
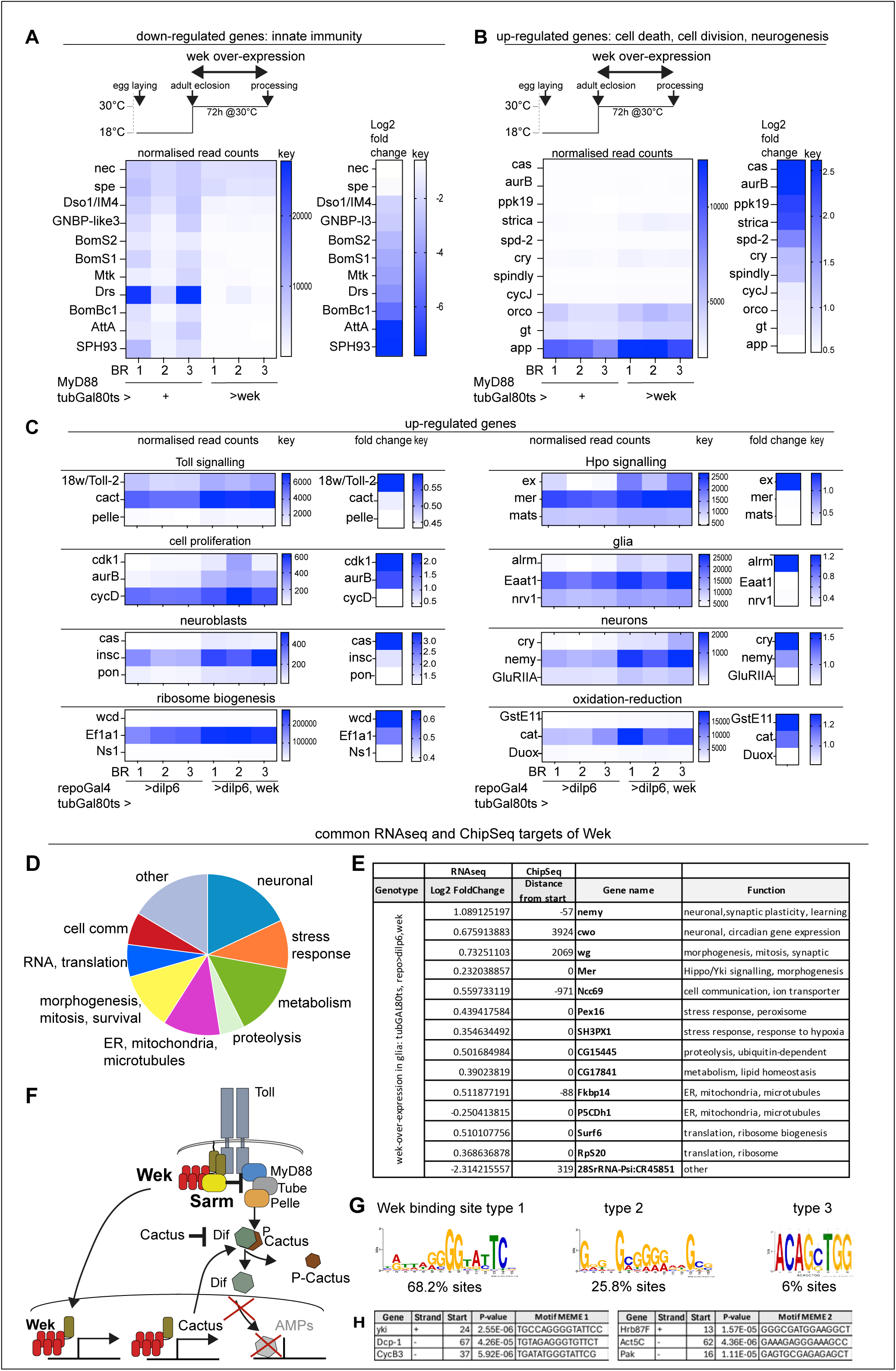
Wek inhibits innate immunity and promotes a regenerative programme. **(A,B)** Bulk RNAseq transcriptomic analysis from fly heads after over-expressing *wek-HA* in MyD88+ cells in adult flies *(tubGAL80^ts^, MyD88>wekHA* vs. *tubGAL80^ts^, MyD88>+* control). See Supplementary Figure S2A. **(A)** Expression of anti-microbial peptide encoding genes and other innate immunity genes was downregulated (see Table S2). **(B)** Expression from genes promoting cell proliferation and neuronal functions was up-regulated (see Table S3). **(C)** RNAseq analysis from fly heads over-expressing *dilp-6* and *wek-HA* in glia, in adult flies *(tubGAL80^ts^, repo>dilp-6, wek* vs *tubGAL80^ts^, repo>dilp-6* controls). This resulted in the up-regulation of the inhibitor of Toll signalling *cactus*, and genes involved in morphogenetic processes. See Supplementary Figure S2B and Tables S4, S5. **(D,E)** Common downstream targets of Wek identified by ChipSeq and RNAseq from adult fly heads over-expressing *dilp-6* and *wek-HA* in glia, in the adult *(tubGAL80^ts^, repo>dilp-6, wek)*. Selection from data shown in Supplementary Figure S3, Tables S6 and Table S7**. (F)** Illustration showing that Wek regulates the expression of genes unrelated to innate immunity and can block innate immunity by linking Tolls to Sarm and by up-regulating the expression of *cactus*. **(G)** ChipSeq identification of three distinct Wek-binding motifs in target genes (see Table S8). **(H)** Examples of Wek target genes involved in cell death, cell proliferation, cell growth and synaptogenesis.

The inhibition of innate immunity by Wek could be due to its ability to bring Sarm – the universal inhibitor of MyD88 - close to MyD88 (Carty et al., 2006; Foldi et al., 2017; Peng et al., 2010). *Sarm* is broadly expressed in neurons, where it drives apoptosis and neurite destruction (Foldi et al., 2017; Karnik and Joshi, 2025; Lin et al., 2014; Mukherjee et al., 2015; Osterloh et al., 2012; Singh et al., 2025). Thus, to investigate further whether *wek* over-expression could switch on alternative cellular programmes, we reasoned that bypassing Sarm would reveal pathways downstream of Wek other than inhibiting immunity and promoting degeneration. MyD88+ cells in the brain include neurons, glial cells and Dpn+ progenitor cells, some of which also express the glial marker *repo* (Li et al., 2020). Toll-2 can induce neurogenesis from MyD88+ Dpn+ Repo+ progenitor cells (Li et al., 2020). Thus, we over-expressed *wek-HA* in glia (using *repoGAL4*), which might target both glial cells and neural progenitor cells, but not neurons. To facilitate the nuclear translocation of Wek and its function as a transcription factor, we co-expressed insulin-like peptide *dilp-6*, and we used *tubGAL80^ts^* to conditionally switch on GAL4 after adult eclosion. Over-expression of *wek* with *dilp-6* up-regulated the levels of *cactus* (Figure 5C, Table S4), the inhibitor of NFκB/Dif/Dorsal. In this way too, Wek repressed canonical Toll signalling, innate immunity and cell quiescence. Furthermore, there was a predominant down-regulation in cell-to-cell communication (Supplementary Figure S2, Table S5) and up-regulation in redox, lipid metabolism, cell proliferation, ribosome biogenesis and translation, neuroblast, neuronal and glial fate genes (Figure 5C, Supplementary Figure S2, Table S4) – altogether, a regenerative signature. These findings suggested that the signalling adaptor Wek could drive cell proliferation and cell growth downstream of Tolls.

Wek could also inhibit Toll-dependent immune signalling by directly repressing the expression of antimicrobial peptides. To test if this was the case and identify direct target genes regulated by Wek, we carried out chromatin immuno-precipitation (ChIP). We over-expressed *wek* and *dilp-6* in adult glia (with *repoGAL4 tubGAL80ts*), precipitated Wek-HA from heads and carried out high-throughput sequencing of bound DNA. Most notably, none of the anti-microbial genes were found to be direct targets of Wek (Supplementary Figure S3, Table S6). Instead, ChipSeq identified as Wek gene targets (Figure 5D, Supplementary Figure S3, Table S6): *wek* itself, meaning that Wek can regulate its own expression; genes involved in cell death (e.g. *Dcp1*), autophagy, neuronal development and function (e.g. *per*), lipid metabolism (eg *mondo*), cell cycle (e.g. *cycB3, Bub3, pbl*), stem cell proliferation (eg *GEFmeso, wdp, Sox15*) and translation (e.g. *ribosomal proteins, translation factors*). Notable targets include genes of the Hippo signalling pathway (e.g. *mer, Tgi* and *yki)* and insulin signalling (eg *PTEN)*. Common Wek target genes identified in both RNAseq and Chipseq included those involved in morphogenesis and regeneration, including translation, stress responses, cell communication, neuronal functions, metabolism, mitochondria, endoplasmic reticulum and microtubules (Figure 5E, Table S7).

ChIPseq analysis revealed three types of Wek binding sequences (Figure 5G, Table S8, Table S9): type 1 of the NFκB class, and shared binding site for Dorsal; type 2 of the SP1 class, shared with buttonhead, required for head formation; and type 3 of the E-box family, shared with Worniu which directs neural stem cell fate. Wek binding motifs were found in the promoter regions of Yki, Dcp1 (apoptosis) and CycB3 (cell division), Act5 (cytoskeleton), Pak (cytoskeleton) and Hrb87F (RNA-binding), amongst others (Table S8, Table S9). Intriguingly, multiple Wek targets identified with ChIPSeq are also Yki targets identified with DamID and ChipSeq, including *wek, cycB3, mondo, Tgi, PTEN, mer* and *Dcp-1* (Oh et al., 2013; Zhang et al., 2017). Like Wek, Yki had also been previously shown to up-regulate the expression of *cactus,* inhibiting innate immunity (Liu et al., 2016). Remarkably, ChipSeq analysis identified *yki* as a direct target of Wek; and *wek* is a target of *yki* (Oh et al., 2013; Zhang et al., 2017).

To conclude, no anti-microbial genes were found to be targets of Wek (Figure 5F). Instead, these data showed that Wek switches Toll signalling from regulating innate immunity and cell death to promoting an alternative morphogenetic or regenerative programme driving cell proliferation, cell growth and cell shape.

### Wek promotes glial cell proliferation and growth together with Yki during brain development

To test whether Wek could regulate glial cell proliferation and growth in vivo, we focused on pupal glioblasts labelled by the reporter *NP6520Gal4*. Neuropile glia in *Drosophila* comprise ensheathing and astrocyte-like glia, which in larvae originate from embryonic glioblasts. During pupariation, larval astrocyte-like glia die, and ensheathing glia become glioblasts that give rise to both astrocyte-like and ensheathing glia, including those labelled by *NP6520Gal4* (Kato et al., 2020). Subsequently, this reporter is switched off from astrocytes to label only adult ensheathing glia (Kato et al., 2020). To ask whether *wek* could regulate glial proliferation, we first used the mitotic marker anti-phospho-Histone-H3 and looked at 8h after puparium formation (APF), when these glial progenitor cells divide (Kato et al., 2020). In fact, we found NP6520>hisYFP+ cells that also had anti-pH3, confirming they were dividing. Over-expression of *wek* resulted in larger hisYFP+ glial cells which were presumably growing, at 8-24h APF there were also pH3+ YFP+ dividing glia, and by 24h His-YFP+ cell number had increased, showing that cells had divided during this period (Supplementary Figure S4). As proliferation was protracted over time, we thereafter quantified cell number in adult brains and counted NP6520>hisYFP+ cells automatically with DeadEasy. Over-expression of *wek* increased the number of NP6520>hisYFP+ glial cells in the brain (Figure 6A,B). By contrast, *wek ^TRiP.GLV21045^-RNAi* caused a reduction in NP6520>hisYFP+ cell number compared to controls (Figure 6A,B). These data showed that Wek can promote and is required for glial proliferation during pupal to adult development. By contrast, over-expression of wild-type *yki* did not affect glial cell number (Figure 6A,B), which could be explained as Yki alone cannot regulate gene expression. Remarkably, over-expression of the Hippo-pathway constitutively active form of Yki *(yki^S168A^)* did not increase glial cell number, and instead reduced it relative to the controls (Figure 6A,B). RNAi knock-down of *yki* had no effect and knocking down both *wek* and *yki (NP6520>hisYFP+, wek ^TRiP.GLV21045^-RNAi, yki ^TRiP.JF03119^-RNAi)* reproduced the effect of knocking down *wek* alone. This could suggest that Wek may function alone rather than with Yki to regulate glial cell number. To test this, we over-expressed both *wek* and *yki* together *(NP6520>hisYFP+, wek, yki)*, and this prevented the upregulation of glial cell number that *wek* alone would have caused (Figure 6A,B).

**Figure 6.**
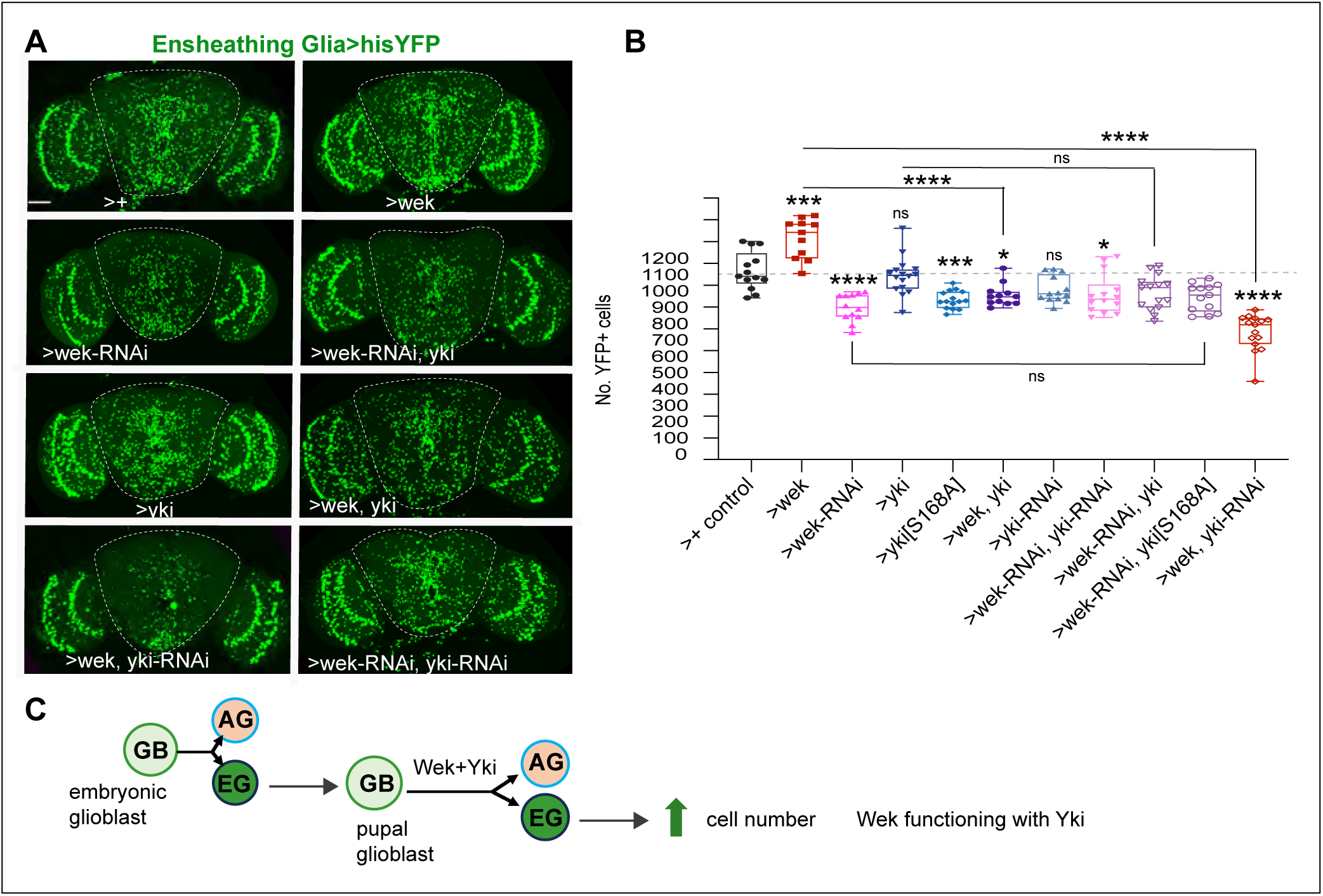
Wek promotes proliferation of glial cells together with Yki. **(A, B)** Over-expression of *wek-HA* in *NP6520>hisYFP+* progenitor cells increase cell number, whereas *wek-RNAi^TRiP.JF03119^* knock-down decreased it, quantification with DeadEasy given in **(B).** Over-expression of either wild-type *yki-V5* or the Hippo-regulated constitutively active *Yki^S168A^-V5* did not increase *NP6520>hisYFP+* glial cell number. Over-expression of both *wek-HA* and *yki-V5* prevented the increase in cell number that *wek-HA* alone would have caused. Genetic epistasis evidence that Wek and Yki function together to promote glial cell proliferation: *yki-RNAi^TRiP.JF03119^* knockdown rescued the increase in *NP6520>hisYFP+* cell number that *wek-HA* over-expression would have caused. Dashed lines indicate ROI used for automatic cell counting with DeadEasy. **(C)** Illustration of the glioblast lineage visualised by NP6520Gal4 in pupae. **(B)** Group One-Way ANOVA, p<0.0001, followed by multiple comparisons Tukey’s tests.*p<0.05, **p<0.01, ***p<0.001, ****p<0.0001. For genotypes and sample sizes, see Table S11. **(A)** Scale bar: 50μm.

We had shown that excess of Wek impaired the nuclear localisation of Yki (Figure 4D), so perhaps excess Wek and Yki compromised their functions. So next, we used genetic epistasis, to test whether *wek* and *yki* might depend on each other or not to regulate glial cell proliferation. We first knocked down *wek* with RNAi together with over-expressing *yki (NP6520>hisYFP, wek-RNAi^TRiP.GLV21045^, yki),* and this resulted in a comparable decrease in glial cell number to that caused by *wek-RNAi^TRiP.GLV21045^* knock-down alone (Figure 6A,B). By contrast, knocking-down *yki* concomitantly with over-expressing *wek (NP6520>hisYFP, wek, yki-RNAi^TRiP.JF03119^)* reduced cell number and prevented the increase in cell number that *wek* over-expression would have caused (Figure 6A,B). Altogether, these data showed that Wek regulates proliferation of ensheathing glia progenitor cells and that Wek depends on Yki to promote glial cell proliferation (Figure 6C).

To ask whether Wek – either alone or with Yki - could regulate glial growth or differentiation, we used the ensheathing glia driver *NP6520GAL4* to express the cell membrane reporter *mCD8-GFP* and measured glial enwrapment of the antennal lobe in adult brains. We found that over-expression of either *wek* or *yki* alone, and also *yki^S168A^*, increased glial membranes (Figure 7A,D). By contrast, neither *wek-RNAi^TRiP.GLV21045^* nor *yki-RNAi^TRiP.JF03119^* knock-down alone had any effect (Figure 7B,C,F). This could be explained because we although *wek-RNAi* knock-down reduced cell number, the remaining cells might compensate for the missing cells by growing larger. To test whether Wek and Yki might function together, we knocked-down *wek* whilst also over-expressing *yki (NP6520>mCD8GFP, wek-RNAi^TRiP.GLV21045^, yki),* and in the absence of Wek, Yki was no longer able to promote glial cell growth (Figure 7B,E). Similarly, we over-expressed *wek* and knocked-down *yki (NP6520>mCD8-GFP, wek, yki-RNAi^TRiP.JF03119^)*, and in absence of Yki, Wek was unable to promote glial growth (Figure 7C,F). These data showed that Yki and Wek function together to promote glial cell growth (Figure 7G,H).

**Figure 7.**
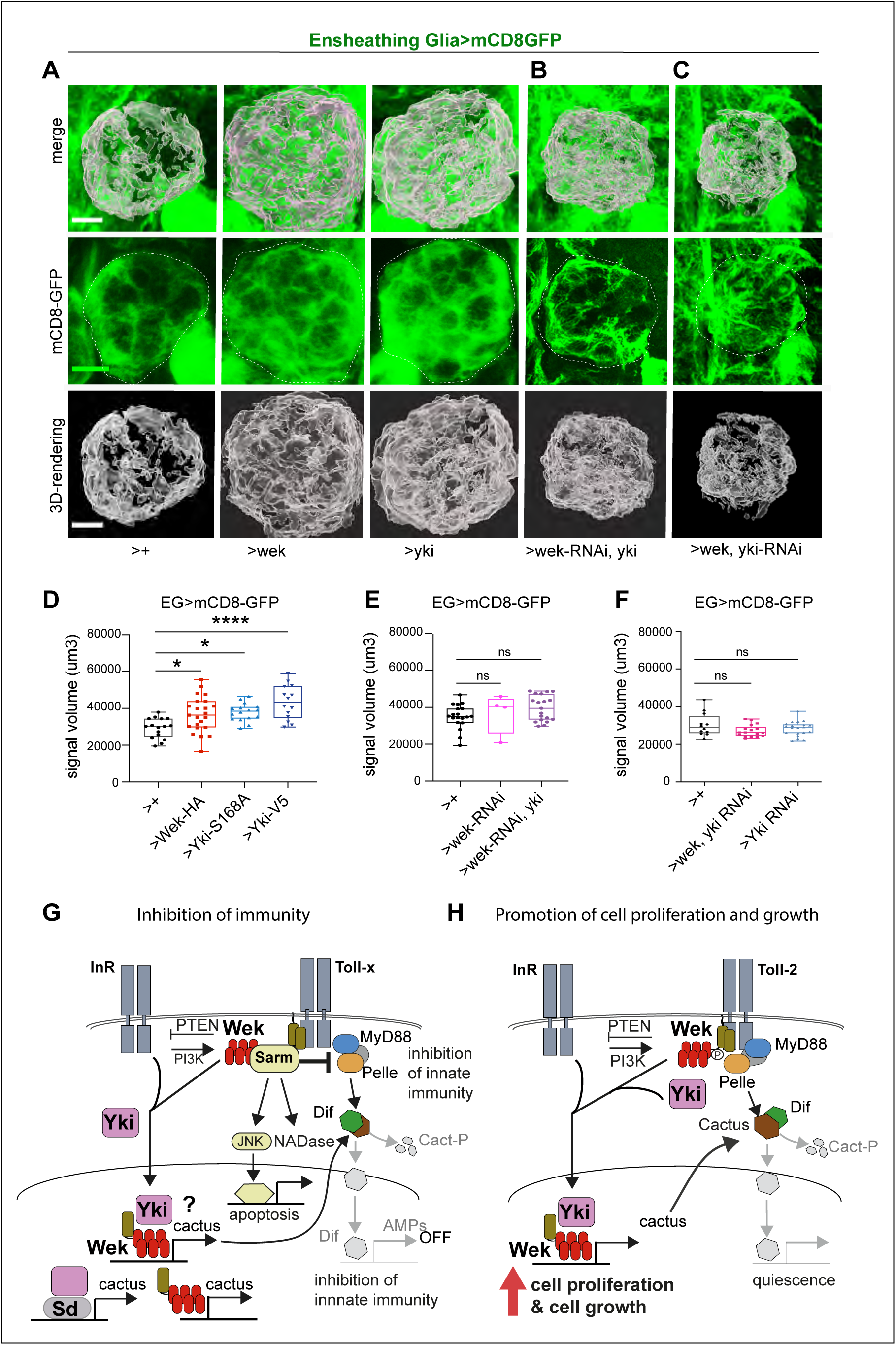
Wek promotes glial membrane growth together with Yki. **(A,D)** Ensheathing glial projections in the antennal lobe visualised with *NP6520>mCD8-GFP* and 3D-volume measured with Imaris. Over-expression of *wek*-*HA*, wild-type *yki-V5* or Hippo-signalling constitutively active *yki^S168A^*, increased glial size. One Way ANOVA p<0.0001, followed by multiple comparisons Tukey’s test: *p<0.05, ****p<0.0001. **(B,E)** Genetic epistasis analysis showing that Wek and Yki together regulate glial growth, as *wek-RNAi^TRiP.JF03119^* knock down rescued the increase in glial growth caused by *yki-V5* over-expression. One Way ANOVA n.s., p=0.136. **(C,F)** Genetic epistasis analysis showing that *yki-RNAi^TRiP.JF03119^* knock down rescued the increase in glial cell growth caused by *wek-HA* over-expression. One Way ANOVA n.s. p=0.1448. **(G)** Summary illustration of how Wek can inhibit innate immunity. At the plasma membrane, Wek links Tolls to Sarm, enabling Sarm to inhibit MyD88 and to promote cell death. Nuclear Wek can upregulate the expression of cactus, which prevents the nuclear translocation of Dif and the expression of AMP genes. Yki functioning with Sd can also up-regulate *cactus* (Liu et al 2016). **(H)** Illustrative summary showing that Wek initially binds Toll-2 and Toll-6 at the plasma membrane, through myristoylation and phosphorylation by Pelle. Insulin or PI3Kinase signalling drive the nuclear translocation of Wek, which carries Yki with it. Wek and Yki promote the expression of genes involved in cell proliferation and growth. The up-regulation of *cactus* expression by Wek releases MyD88-dependent cell quiescence. **(A-C)** Dashed lines indicate antennal lob ROI measured in 3D with Imaris. Scale bars: 50μm. See Table S11 for genotype details and sample sizes.

Altogether, these data show that Wek and Yki function together in vivo during development of the adult brain, to promote glial cell proliferation and growth.

### *wek* orthologues in humans?

To seek potential *wek* orthologues in humans, we carried out a bioinformatics analysis. A PhylomeDB search using Wek as query identified six putative human orthologs of *Drosophila wek*, three of which were supported with high confidence (consistency score =1) (Table S10). InterPro analysis further revealed five human proteins containing a ZAD domain, one of which was invalid and the other four were all encoded by the same gene, *ZFP276*. We subjected all seven genes to domain analysis using InterPro and confirmed conserved zinc-finger domains across all, but only one human ZAD-containing protein (Q8N554) encoded by the *ZFP276* gene (Supplementary Figure S5). ZFP276 lacks a myristoylation consensus sequence.

Notably, Q8N554 exhibits a similar domain organisation to that of *Drosophila* Wek, but PhylomeDB did not identify *ZFP276* as a direct orthologue of *wek* (Table S6). This suggests that the ZAD domain could have been acquired independently in different lineages, leading to convergence in domain organisation without underlying orthology. Instead, putative ZFP *wek* orthologues in humans contained the related BTB, as well as SAM-binding domains (Table S10, Supplementary Figure S5).

## DISCUSSION

Toll and Toll-Like Receptors (TLRs) are best known for their universal function promoting innate immunity and neuroinflammation (Hoffmann, 2003; Imler and Hoffmann, 2001; Kumar, 2019; Pascual et al., 2021; Squillace and Salvemini, 2022). However, *Drosophila* Tolls and mammalian TLRs also have non-immune functions in many tissues, including the brain. These include regulating cell-cell interactions, cell survival and death, cell proliferation (neurogenesis, gliogenesis), cell size (neurite growth and retraction), structural synaptic plasticity and behaviour (Anthoney et al., 2018; Chen et al., 2019; Church et al., 2016; Foldi et al., 2017; Hung et al., 2018; Li et al., 2020; Li et al., 2021; McIlroy et al., 2013; Okun et al., 2010; Okun et al., 2011; Pare et al., 2014; Rolls et al., 2007; Sanchez-Petidier et al., 2022; Shechter et al., 2008; Shen et al., 2016; Sun et al., 2024; Tamada et al., 2021; Zhang et al., 2024). Some of these cellular features are regulated during structural brain plasticity, which underlies learning, adaptation and anti-depressant action (Troubat et al., 2021). By contrast, neuro-inflammation is emerging as a key driver of ageing, neurodegenerative and psychiatric diseases (Troubat et al., 2021). The ability of Tolls and TLRs to induce both regenerative and degenerative programmes is key to revealing how the brain can switch from a plastic to an inflammatory state. Here, we have shown that Wek is a Toll signalling adaptor that inhibits innate immunity and promotes a morphogenetic or regenerative programme instead.

Toll/TLR receptors bind signalling adaptors through interactions between TIR domains present in both (Watters et al., 2007). Instead, Wek is a ZAD-ZFP lacking a TIR domain that directly binds the TIR domains of at least Toll-2 and Toll-6. Wek is associated to the plasma membrane by myristoylation, where it is phosphorylated by the Toll signalling kinase Pelle. Excess of Pelle and Wek drove nuclear degradation, consistent with the function of Wek binding Sarm to drive cell death downstream of Tolls (Foldi et al., 2017; Singh et al., 2025). Phosphorylation by Pelle also enables further phosphorylation events that in concert with insulin and/or PI3Kinase signalling drive the nuclear translocation of Wek. Within nuclei, Wek functions as a transcription factor, at least partly together with Yki. We have shown that Wek and Yki could physically interact in co-immunoprecipitation, pull-down assays and in vivo; the nuclear translocation of Wek depends on concerted Toll and insulin/PI3kinase signalling, and Wek enabled the nuclear translocation of Yki; and using genetic epistasis analysis, we showed that Wek and Yki function together in vivo to promote glial proliferation and growth.

Wek can inhibit innate immunity in multiple ways (Figure 7G). In S2 cells, excess of Pelle and Wek drove nuclear degradation, consistent with the function of Wek binding Sarm to drive cell death downstream of Tolls (Foldi et al., 2017; Singh et al., 2025). At the membrane Wek can link Tolls to Sarm, which inhibits signalling downstream of MyD88 and leads to immune evasion (Foldi et al., 2017; Singh et al., 2025). Functioning as a transcription factor, Wek caused a pervasive downregulation of genes involved in innate immunity, prominently AMP gene expression. However, it did not directly regulate these genes. Instead, Wek suppresses innate immunity indirectly, by facilitating the inhibition of MyD88 by Sarm downstream of Tolls, and by up-regulating the expression of *cactus*. In fact, Wek-pull down brought down Sarm, consistent with previous evidence that Wek can physically interact with Sarm downstream of Tolls (Foldi et al., 2017). *wek* over-expression raised the levels of *cactus*, the inhibitor of NFκB/Dif, which would block an immune response. Yki has been previously shown to inhibit innate immunity in this way too (Liu et al., 2016). Yki is an effector of Hippo signalling and its nuclear exclusion is driven by its phosphorylation downstream of Hippo or AMPK signalling (Davis and Tapon, 2019; Gailite et al., 2015; Huang et al., 2005; Zheng and Pan, 2019). Yki is a co-activator that regulates gene expression by interacting with transcription factors, most commonly Scalloped (Sd) (Davis and Tapon, 2019; Goulev et al., 2008; Zheng and Pan, 2019). Nuclear Yki with Sd up-regulates *cactus* expression, which prevents Dif/NFκB from entering the nucleus (Liu et al., 2016). However, upon infection, Toll-1 activates Pelle, which phosphorylates the Hippo inhibitor Cka, promoting Hippo signalling and Yki nuclear exclusion, thus enabling Dif/NFκB to drive immunity (Lemaitre et al., 1996; Liu et al., 2016). Interestingly, Wek also up-regulates *cactus* expression, suggesting that Wek and Yki could induce *cactus* expression either together or synergistically, further inhibiting the immune response. Thus, in the brain Toll-1 activation can drive both innate immunity and the degenerative Wek-Sarm routes downstream. Remarkably, activation of Toll-1 during innate immunity inhibits release of insulin-like peptides and signalling downstream of PI3Kinase in the fat body (Roth et al., 2018; Suzawa et al., 2019). This would prevent the nuclear translocation of Wek, and of Yki with it. In this way, infection diverts Toll-1 signalling towards innate immunity (via MyD88) and degeneration (via Sarm), and away from the morphogenetic or regenerative Wek-Yki route.

By contrast, in the absence of infection, our findings link Wek to Yki directly downstream of Toll signalling to promote cell proliferation and cell growth (Figure 7H). We showed that the nuclear translocation of Wek depends on concerted Insulin Receptor and Toll signalling, critically involving PI3Kinase. Furthermore, in the fat body and in the absence of infection, the nuclear translocation of Yki depends on Wek and Pelle. When *pelle* was knocked-down, Yki was excluded from nuclei. Sustained *pelle* over-expression reduced Yki’s nuclear distribution over time. The latter was consistent with findings that over-expression of *pelle* phosphorylates Cka, enabling Hippo signalling, to promote the cytoplasmic retention of Yki (Liu et al., 2016). The phosphorylation of Wek - and potentially Yki - by Pelle might help them shuttle to the nucleus together: (1) the phosphorylation of Wek at Ser164 enabled further phosphorylation at Ser168. (2) PI3Kinase drove phosphorylation of Wek at Ser168, and mutant Wek^S168A^ prevented the nuclear translocation of Yki in vivo. (3) Precipitation of Yki from the nuclear cellular fraction brought down Wek phosphorylated at both Ser 164 and Ser168. Together, these data suggest that Wek phosphorylation at Ser168 is required for the nuclear translocation of both Wek and Yki, and for the interaction between Wek and Yki. Perhaps Pelle facilitates the interaction between Wek and Yki at the plasma membrane, enabling their subsequent nuclear translocation. Importantly, these data indicate that Yki is a downstream target of Tolls via the non-canonical Wek signalling pathway.

Identifying which Tolls might drive concerted Wek and Yki nuclear shuttling was beyond the scope of this work, but Toll-2 is a prime candidate. Tolls are not all equal, and infection results in the activation of Toll-1 and Toll-5, but not Toll-2 or Toll-6 (Tauszig et al., 2000). Furthermore, Toll-2 over-expression induced cell proliferation in the adult by inhibiting MyD88-dependent cell quiescence (Li et al., 2020); Toll-2 induced adult neurogenesis depends on both *wek* and *yki* (Li et al., 2020); and Toll-2 can directly link to PI3Kinase (Tamada et al., 2021). Thus, at least Toll-2 and Toll-6 are ideally placed to promote cell growth via Wek.

Our data show that Wek switches Toll signalling from innate immunity towards a gene expression signature characteristic of morphogenetic and regenerative processes. Wek can regulate gene expression independently of Yki, and also together with Yki, presumably by targeting distinct genes. In fact, Wek (this work) and Yki (Kwon et al., 2015; Oh et al., 2013; Zhang et al., 2017) RNAseq and DNA-binding (ChipSeq and DamID) gene targets overlapped but also differed, although so did the cells of origin for these studies. Our in vivo data showed that Wek regulates glial cell proliferation and growth together with Yki. In this context, over-expressed wild-type Yki or Hippo-pathway activated Yki^S168A^ alone could induce glial cell growth but not glial cell proliferation in the central brain. This suggests that Yki and Yki^S168A^ might also regulate distinct genes from Wek controlling cell size control in the central brain. This could mean that Wek and Yki together regulate genes involved in both cell growth and cell proliferation, and Yki potentially with Sd might also regulate another gene subset involved in glial cell growth. On the other hand, both Yki and Sd have been previously found to regulate glial cell proliferation in optic lobes (Reddy and Irvine, 2011). Future work could determine whether perhaps Wek, Sd and Yki form ternary complexes to regulate expression of shared target genes. Alternatively, Wek and Sd may compete for binding to Yki. Or distinct cell types may express *wek* or *sd* or other *yki* partners such as *hth*, driving unique cell-type specific interactions with Yki.

How could Wek, a Toll signalling adaptor, switch the outcome of Toll signalling from innate immunity and cell death to activating a generative programme instead? Three factors could be involved: (1) whether Sarm is present in the cell or not. In the presence of Sarm, Wek can promote cell death downstream of Toll-1, Toll-6 and Toll-7 and immune evasion downstream of Toll-1 (Foldi et al., 2017; Singh et al., 2025). Sarm is the universal inhibitor of MyD88, it is expressed in neurons throughout animals (Lin et al., 2014; Mukherjee et al., 2015; Osterloh et al., 2012), but perhaps not in other cell types. (2) The specific Toll that Wek interacts with in each context could drive distinct outcomes, as the functions of each Toll receptor differ (Chauhan et al., 2024; Foldi et al., 2017; Lemaitre et al., 1996; Li et al., 2020; McIlroy et al., 2013; Nakamoto et al., 2012; Narbonne-Reveau et al., 2011; Singh et al., 2025; Tauszig et al., 2000)). (3) Insulin or PI3K signalling will influence the response, as they are required for Yki activation (Strassburger et al., 2012) and nuclear translocation of Wek. Altogether, our data indicate that the switch from immune or degenerative responses to a generative response involves concerted Toll-2 or Toll-6 and insulin/PI3Kinase signalling to engage Wek and Yki.

Why does the shared function of Toll receptors, Wek and Yki matter? Yki is a critical inducer of cell survival, cell proliferation, neural stem cell activation and cell growth (Davis and Tapon, 2019; Ding et al., 2016; Ding et al., 2020; Manning et al., 2020; Mo et al., 2014; Zheng and Pan, 2019) . These cellular processes underlie development, cell competition, regeneration and cancer, and Yki is involved in all of these (Ding et al., 2020; Zheng and Pan, 2019). The induction of neurogenesis, gliogenesis and cell growth in adult brains are key manifestation of brain plasticity, induced by learning, exercise and anti-depressants (Troubat et al., 2021). Thus, a link between Tolls/TLRs, Wek and Yki could have important implications on how these receptors can switch away from innate immunity and an inflammatory state to controlling cell proliferation and cell growth instead. In mammals, the signalling adaptor B-cell activator protein (BCAP) also regulates two distinct signalling pathways. BCAP has an N-terminal TIR domain that interacts with adaptors MyD88 and Mal to modulate inflammatory signalling by TLR2 and 4 (Troutman et al., 2012a). It is also required for the activation of signal transduction by the B-cell receptor (BCR). In particular, BCAP couples the BCR to the activation of PI3 kinase which drives B-cell proliferation through the protein kinase Akt1 (Lauenstein et al., 2019; Okada et al., 2000; Troutman et al., 2012a; Troutman et al., 2012b). Thus, polyfunctional signalling by Tolls and TLRs may be conserved during evolution. In humans, there is only one ZAD-ZFP, ZFP276, and it can inhibit oligodendrocyte progenitor cell proliferation and promote oligodendrocyte differentiation, although findings are extremely limited (Aberle et al., 2022; Chung et al., 2007; Wong et al., 2003). Similarly to ZFP276 (Aberle et al., 2022), we have shown that Wek also promotes glial cell growth. However, our bioinformatics analysis revealed that *ZFP276* is not a *wek* orthologue, and instead O15060 encoded by *ZBTB39* containing a BTB instead of a ZAD domain appears to be more closely related. Intriguingly, *wek* candidate orhologues Q9NQX0 and A0A024RBM7 contain SAM domains, which could interact with Sarm. However, ZFPs duplicated multiple times throughout animal lineages and it is not possible to assign accurate orthology relationships to *wek*. ZAD-ZFPs are most prevalent in arthropods, but, ZAD, BTB, SCAN and KRAB ZFPs evolved for genome defence and co-evolved into species-specific forms with the transposons they tamed (Angileri et al., 2022; Chung et al., 2007; Church et al., 2016; Collins et al., 2001; Ecco et al., 2017; Edelstein and Collins, 2005). Importantly, these kind of ZFPs are generally involved in the control of stemness and cell fate. Thus, a ZFP containing a BTB, SCAN or KRAB domain could play the function of Wek downstream of TLRs in humans. Intriguingly, TLR4 promotes OPC proliferation and differentiation (Church et al., 2016), and it is compelling to test whether a functional relationship between TLR4 and ZFPs exists.

To conclude, we have shown that Wek is a molecular switch that can drive Toll signalling away from innate immunity and towards a morphogenetic or regenerative programme instead. This requires concerted insulin or PI3K and Toll signalling, enabling Wek to function together with Yki to promote cell proliferation and cell growth. We provided evidence that this mechanism operates during brain development. However, the findings suggest the same mechanism could drive cell proliferation and growth downstream of Tolls and TLRs also during adult structural brain plasticity, and outside the brain, with broad important implications for contexts such as cell competition, regeneration and cancer.

## MATERIALS AND METHODS

***Drosophila* genetics.** Please see Table S11 for details on experimental genotypes and sample sizes and Table S12 for the list of the stocks used. Stocks carrying combinations of multiple drivers, responders and other tools were generated by conventional genetics. To identify balancer chromosomes in larvae, we used *SM6aTM6B, Tb-*, with which the second and third chromosome co-segregate. For adult brains in Figures 6 and 7, only females were used. **Knock-in-allele:** *w; wek-HA^CRISPR-KI^* (this work) is an endogenously tagged allele, at the C-terminus (see below); **GAL4 driver lines:** *ppl-GAL4* drives expression in the fat body; *repoGal4* drives expression in all glial cells; NP6520Gal4, drives expression in ensheathing glia. **UAS lines:** (1) **for knock-down or down-regulation:** UAS-MyD88 RNAi^GD9716^ (VDRC 25399); UAS-*pelle-RNAi^TRiP^ ^HMS04458^*; *UAS-PTEN-H* (BSC 82170); *UAS-InR-DN^K1409A^* (BSC 8252); *UAS-wekRNAi^TRIP^ ^GLV21045^@attP2* (BSC 35680); *UAS- wek RNAi^TRIPMHC04653^@attP40* (BSC 57260); *UAS-yki-RNAi^TRIPF03119^@attP2* (BSC 31965); (2) **for over-expression:** UAS-wek-HA (FlyORF), *UAS-wek-FLAG* (this work), *UAS-wek^S168A^-FLAG* (this work); *UAS-dilp-6* (gift of E. Hafen); *UAS-yki-FLAG* (BSC 600529); *UAS-yki-V5* (BSC 28819); *UAS-yki^S168A^-V5* (BSC 28818); *UAS-mCD8-GFP*, membrane tethered Green Fluorescent Protein; *UAS-histone-YFP*, nuclear Yellow Fluorescent Protein. **Conditional expression** was carried out by combining Gal4 driver lines with using *tubGAL80^ts^*, which expresses the Gal4 inhibitor at 18°C but it is not functional at 30°C. For RNAseq and ChipSeq (Figure 5), flies were kept at 18°C until adult eclosion and then shifted to 30°C for 3 days (see below). For larval fat body analysis (Figure 4F), larvae were kept at 18°C from egg laying for 8 days (192h) and then shifted to 30°C for 20h or 48h and dissected at wandering stage (*tubGal80ts, ppl>Yki-V5* (Control) and *tubGal80ts, ppl>Yki-V5, Pelle-HA)*.

### Bioinformatics

We investigated the evolutionary relationships of the *Drosophila melanogaster wek* gene family with potential human orthologs. Candidate orthologs were identified using PhylomeDB (https://doi.org/10.1093/nar/gkab966), which provides precomputed gene trees and orthology predictions across multiple species. Orthology relationships were classified according to the PhylomeDB consistency score and support across different phylomes. Protein domain architectures of the candidate orthologs were annotated using InterPro (https://doi.org/10.1093/nar/gkae1082). We examined both the predicted human orthologs of *wek* and human proteins containing ZAD motifs. Domain organisation plots were generated directly from InterPro annotations.

### Molecular Biology

#### Molecular cloning and Site-Directed Mutagenesis

See Table S13 for the list of constructs and primers used for molecular cloning. To express *wek-ZAD, wek- FL, Toll-2ICD* and *Toll-6ICD* for protein purification, the corresponding DNA inserts were cloned into pE15b plasmid for bacterial expression (please see below under “Protein expression and purification”). To over-express wild-type *wek-FLAG* in S2 cells and in vivo, *wek* coding cDNA was PCR amplified from BDGP gold clone LD22579 using Phusion High-Fidelity DNA polymerase (New England Biolabs), following Gateway cloning procedures inserting them first into *pDONR* and subsequently into *pAct5c-3xFlag* (Table S13). Point mutations *(pAct5c wek-FLAG, pAct5c wek-G2AFLAG, pAct5c wekS6N-FLAG, pAct5c wekS164AFLAG, pAct5c wekS168AFLAG)* were generated into *pAct5c-Wek-Flag WT* (wild type) using Q5 ® Site-Directed Mutagenesis (NEB, cat. Number E0554S) following manufactureŕs instructions, by inserting the desired mutations – *wekG2A, wekS6N, wekS164A, wekS168A* – into primer sequences (Table S13) for cloning into *pAct5. UAS-wek-FLAG* and *UAS-wekS168A-FLAG* were generated by Gateway Cloning, inserting the coding regions into pDNOR, see Table S13 for primers. Transgenesis was subcontracted to FlyORF and *w+* stable stocks were generated and balanced.

Flies bearing an endogenous HA tag at the C-terminus of *wek* (i.e. *wek-H^ACRISPR-KI^* allele) were generated by Crispr-Cas9 enhanced homologous recombination and transformant flies were identified by 3xP3-RFP which as flanked by LoxP sites (outsourced to WellGenetics), and subsequently removed with CRE-recombinase. Final stocks were established from single males followed by balancing.

#### Bulk RNA sequencing (RNAseq) from *Drosophila* brains

Total RNA was extracted from *Drosophila* melanogaster heads using the TRIzol method, followed by column purification. For each biological replicate, 50–100 heads were pooled and transferred to 1.5 ml RNase-free tubes containing 500 µl of TRIzol reagent (Thermo Fisher Scientific), after which they were immediately frozen at -80 °C. The tissue was then homogenised using a micro-pestle and incubated at room temperature for five minutes. To induce phase separation, 100 μL of chloroform was added to each sample. The samples were then vigorously vortexed for 15 seconds, incubated at room temperature for 10 minutes, and centrifuged at 12,000×g for 15 minutes at 4 °C. The RNA was then precipitated from the aqueous phase by adding an equal volume of isopropanol. The mixture was incubated at -20°C overnight, after which it was centrifuged at 12,000 × g for 10 minutes at 4°C. The supernatant was then discarded, the RNA pellet washed with 500 μL of 75% ethanol by centrifugation at 7,500×g for 5 minutes at 4 °C and air-dried for 5– 10 minutes. The pellet was then resuspended in 30–50 μL of RNase-free water, incubated at 55 °C for 5 minutes, and treated with DNase I (Qiagen) to remove DNA contamination. The RNA concentration and purity were assessed using a NanoDrop spectrophotometer (Thermo Fisher Scientific) with an A260/A280 ratio between 1.8 and 2.1, and verified by Novogene. Novogene carried out the sequencing as follows: RNA integrity was verified using an Agilent Bioanalyzer and only samples with an RNA Integrity Number (RIN) greater than 7.5 proceeded to library preparation for RNA sequencing. The cDNA libraries were then sequenced on a NextSeq 500 High Output Flow Cell Set (Illumina) to generate 75-base-pair paired-end reads. Identification of differentially expressed genes was subcontracted to Novogene.

#### Chromatin immune-precipitation and DNA sequencing (ChipSeq)

Tissue samples (20-30 adult fly brains and 20 larva fly brains) were harvested and immediately placed in ice-cold phosphate-buffered saline (PBS) containing 0.1% PBS-T. Cross-linking was performed by incubating the tissue in 1% formaldehyde for 5 minutes at room temperature. The reaction was quenched by adding 125 mM glycine and incubating for 5 minutes at room temperature. The tissue was subsequently centrifuged at 2000 rpm for 5 minutes at 4°C. The resulting pellet was washed twice with ice-cold PBS supplemented with 1x Complete Mini Inhibitor. Pellets were resuspended in 1 mL of cell lysis buffer per 0.5 mL of tissue, consisting of 5 mM PIPES (pH 8.0), 85 mM KCl, 0.5% NP-40, 1x Complete Mini EDTA-free Protease Inhibitor, 1 tablet PhosStop, 1% SDS and samples were incubated on ice for 10 minutes. The buffer was removed, and the pellet was resuspended in nuclease buffer and incubated at 4°C for 10 minutes. The chromatin was diluted in 500 μL IP dilution buffer (16.7 mM Tris pH 8.0, 1.2 mM EDTA, 167 mM NaCl, 1.1% Triton X-100, 0.01% SDS) and sonicated for 5 cycles at 30 s OFF at maximum intensity using a Bioruptor sonicator (Diagenode). Samples were centrifuged at 13000 rpm for 20 minutes in a microcentrifuge at 4°C. The clear supernatant containing fragmented chromatin was transferred to a fresh 15 mL tube, and a 20 μL aliquot was saved for input DNA control. The supernatant was further diluted with 5 vol of IP dilution buffer and the appropriate antibody (5-10 μg) was added to the sample and incubated overnight at 4°C on a rocker. Pre-washed 20 μL Dynabeads were added to the lysate-antibody mix and incubated further for 1 hr at 4°C on a rocker. Beads were pelleted by centrifugation at 3000 rpm, and the buffer was carefully removed and saved for reference. Beads were washed six times with 1 mL of ice-cold low salt buffer containing 0.1% SDS, 1% Triton X-100, 2 mM EDTA, 20 mM Tris pH 8.0, and 150 mM NaCl. A final wash was performed using 2 mL of high salt buffer (0.1% SDS, 1% Triton X-100, 2 mM EDTA, 20 mM Tris pH 5.0, 500 mM NaCl). Beads were washed once with TE buffer (10 mM Tris pH 8.0, 1 mM EDTA).

Chromatin was eluted by incubating beads in a fresh tube with 250 μL of elution buffer (0.1 M NaHCO3, 1% SDS) at room temperature for 15 minutes. Reverse cross-linking was performed by adding 38 μL of de-crosslinking buffer (2 M NaCl, 0.1 M EDTA, 0.4 M Tris pH 7.5) and incubating overnight at 65°C on a rocker. Following overnight incubation, proteins were digested by adding 2 μL of Proteinase K (50 mg/mL) and incubating at 50°C for 2 hours on a rocker. DNA was purified using the Monarch PCR and Cleanup Kit according to the manufacturer’s instructions. DNA concentration and quality were assessed using real-time PCR quantification. For NGS sequencing, ChIP and input DNA were further fragmented to 200 bp fragment size using a Bioruptor Pico (Diagenoide). All ChiP-DNA libraries were produced using the NEBNext Ultra II DNA Library Prep kit (New England Biolab and NEBnext Multiplex Oligos for Illumina Dual Index Primers.

Constructed libraries were assessed for quality sing the Tapestation 2200 with High Sensitivity D1000 DNA ScreenTape. Libraries were tagged with unique barcodes and sequenced simultaneously on a HiSeq4000 sequencer. Library production and sequencing was carried out at the Envision facility, at the University of Birmingham. Bioinformatics analysis of sequences was subcontracted to Marbyt (https://marbyt.com/).

Analysis of hits and identification of GO terms was carried out using Metascape (https://metascape.org/gp/index.html#/main/step1) (Zhou et al., 2019).

### Protein biochemistry

#### AlphaFold models

The full-length Wek protein sequence of *Drosophila* melanogaster was obtained from UniProtKB/Swiss-Prot, accession Q9VJN5. For modelling the ZAD domain, only residues 10-82, as annotated in UniProt, were used. Structural prediction modelling was done using AlphaFold v2.1 1 (Jumper et al., 2021) and the open-source ColabFold (Mirdita et al., 2022).After checking the AlphaFold Protein Structure Database for preexisting models, the query amino-acid sequence was submitted to the prediction pipeline. The pipeline performed multiple sequence alignments (MSAs) and template searches (ColabFold uses MMseq2 for rapid MSA construction), and generated models using the standard AlphaFold2 model ensemble. For each target, five independent predictions were produced and ranked by internal AlphaFold confidence metrics and the confidence of the models was assessed using the per-residue pLDDT score and the predicted aligned error (PAE) between domains. The highest-confidence model was then used for downstream structural analysis and comparison with the known fold.

#### Protein expression and Purification

For recombinant protein production, *wek* was cloned into a modified *pET15b* vector encoding a C-terminal Strep tag preceded by a TEV protease cleavage site. The intracellular domain (ICD) domain of *Toll-2* (synonym *18wheeler*) (residues A1041–L1184) and *Toll-6* (residues Y1079–A1414), and the ZAD domain of *wek* (residues Y10–L85) were amplified by PCR and ligated into *pET15b* vectors digested with NdeI and XhoI. All constructs were verified by Sanger sequencing. Recombinant proteins were expressed in *Escherichia coli* BL21 Rosetta cells (Novagen). **Wek:** Cells expressing *wek-FL* and *wek-ZAD* were grown in LB medium at 310 K with shaking (220 rpm) until reaching an OD600 of 0.8. The temperature was then reduced to 293 K, protein expression was induced with 1 mM IPTG, and cultures were incubated for 12– 16h. Cells were harvested and resuspended in Buffer A (20 mM Tris-HCl pH 8.0, 500 mM NaCl) supplemented with lysozyme and 1 mL BugBuster (Novagen), and lysed using an Emulsiflex C5 homogenizer (Avestin). The lysate was clarified by centrifugation at 14,000 rpm for 30 min at 277 K and loaded onto a 5 mL StrepTrap HP column (GE Healthcare). After washing with 10 column volumes of Buffer A, bound proteins were eluted with Buffer A containing 10 mM desthiobiotin (Sigma). For final purification, proteins were diluted in SEC buffer (20 mM Tris-HCl pH 8.0, 100 mM NaCl), concentrated, and separated on a Superdex 200 10/300 GL column (Cytiva). **Toll-2/-6.** Cells expressing *Toll-2-ICD* and *Toll-6-ICD* were grown in LB medium at 310 K with shaking (220 rpm) until reaching an OD600 of 0.8. Expression was induced with 1 mM IPTG at 298 K and continued for 6 h. Cells were lysed by sonication in lysis buffer (25 mM Tris-HCl pH 7.5, 300 mM NaCl) supplemented with protease inhibitors. The clarified lysate was applied to a His- GraviTrap column (GE Healthcare), and 6xHis-tagged proteins were eluted with a 50–500 mM imidazole gradient. Fractions containing Toll proteins were pooled and desalted on an HP26/10 column (GE Healthcare) into buffer containing 25 mM Tris-HCl pH 7.5, 100 mM NaCl. His-tags were removed by thrombin cleavage, and the proteins were incubated with p-aminobenzamidine resin to remove residual thrombin. Proteins were concentrated using Ultrafree-5 centrifugal filter units (Millipore) and further purified by size-exclusion chromatography on a Superdex 200 10/300 GL column (GE Healthcare) equilibrated in 25 mM Tris-HCl pH 7.5, 100 mM NaCl.

#### Analytical Ultra-Centrifugation

Sedimentation velocity measurements were performed using a Beckman XLA centrifuge at 20 °C in 50 mM Tris, 300 mM NaCl, 1𝜇M ZnCl2, pH 8. Both sample and reference volumes were 400 µl and data were acquired at 50,000 rev/min using an An60Ti rotor (Beckman Coulter). Buffer viscosity and density were estimated using SEDNTERP (Laue et al., 1992) and the data analysed with SEDFIT (Schuck, 2003).

#### Dynamic Light Scattering

Full-length Wek (Wek-FL), the TIR domain of Toll-2 and Toll-6 were column purified following expression in *E.coli* (see above). Wek-FL, Toll2-TIR and Toll6-TIR were passed through 20um filters prior to use. Interactions between proteins were tested using Dynamic Light Scattering (DLS) measurements were performed at 298K on filtered protein samples at 25°C using a Zetasizer Nano ZS (Malvern Panalytical) and a Hellma quartz cuvette with a 3mm path length centered at 9.65mm. 20uL of Wek-FL (22 μM) was placed into the cuvette and aliquots of 44 μM of Toll2-TIR and Toll6-ICD were added so that the total volume in the cuvette did not exceed 26uL. Data were analysed with Zetasizer software (version v8-01, Malvern) using the continuous hydrodynamic radius distribution (Rh, Z-average) model in SEDFIT (Stetefeld et al., 2016)and the polydispersity index (PDI) using the cumulants analysis. In addition, size distributions by intensity, volume, and number were obtained to identify the presence of possible multiple species. Three independent measurements per sample were recorded.

#### Myristoylation detection with click chemistry

To test if Wek is myristoylated, S2 cells were cultured in Schneideŕs *Drosophila* medium (Thermo Fisher Scientific) supplemented with 10% heat-inactivated fetal bovine serum (FBS) and 1% penicillin-streptomycin. Cells (2 x10^5^) were plated into 6-well plates 1 day prior to transfection and grown to 60-70% confluence. Cells were transfected with *pAct-wek-FLAG* and *pAct-WekG2A-FLAG* tagged constructs using 2.5μL of TransIT-Insect (Mirus), according to manufacturer’s instructions. Cells were treated for the experiment after 48 hours. Then they were washed with serum free buffer and incubated for 10 h at 27°C in Complete Media containing 2% v/v GCS and 25 μM Alk-Myr. After the incubation the cells were washed three times with Dulbeccós phosphate-buffered saline (DPBS) and harvested in RIPA buffer (50 mM Tris- HCl, pH 7.4, 150 mM NaCl, 1% NP-40, 0.5% sodium deoxycholate, 0.1%SDS) supplemented with protease inhibitors for 20 min on ice. The samples were then analysed by SDS-PAGE and fluorography. Alk-Myr labelled samples were reacted with Az-TAMRA using a click chemistry reaction cocktail (1 μL Az-TAMRA, 1 μL TCEP, 1 μL TBTA, and 1 μL CuSO4), in a total reaction volume of 50 μL for 1 h at room temperature. After CuAAC, 500 μL of MeOH was added to the sample, then the samples were stored at -80°C overnight. After centrifugation at 1500 rpm at for 30 min at 4°C, the supernatant was removed, and the pellet was washed with 500 μl of MeOH and then dried in air. The samples were denatured by sonication in SDS-sample buffer and then subjected to SDS-PAGE. In-gel fluorescence analysis of the gel obtained by SDS-PAGE was visualized using an Odyssey infrared imaging system.

#### Native gel electrophoresis

Native polyacrylamide gel electrophoresis (Native PAGE) was performed using Bio-Rad NativePAGE™ Novex® 4–16% Bis-Tris gels following the manufacturer’s instructions. Recombinant Wek FL and the TIR and ICD domains of Toll2 and Toll6 were prepared in NativePAGE™ Sample Buffer without reducing agents or SDS to preserve their native conformations. For interaction studies, Wek FL was mixed with Toll2-ICD or Toll6-TIR domains at a 1:1 molar ratio and incubated for 15 minutes at 4°C. As controls, Wek FL and the Toll domains were also loaded separately under identical conditions. Samples (5–20 µg per lane) were applied to the gel, which was run at 150 V for 1.5–2 hours at 4°C in NativePAGE™ Running Buffer. Gels were subsequently stained with Coomassie Brilliant Blue R-250 for 1 hour and destained in 40% methanol and 10% acetic acid until protein bands were clearly visible. Band patterns were compared to the controls to assess complex formation. All steps were performed at low temperature to maintain protein–protein interactions, and no denaturing agents were used, allowing visualization of oligomeric states and complex formation.

#### Pull-down and proteomics

S2 cells were transfected with either *pAct-wek-FLAG* and *Toll-6HA,* or *pAct-wek-FLAG* and *pAct-pelle-HA* and stimulated with DNT-2. S2 cells were grown and transfected as described above. DNT2 protein was expressed and purified from S2 as published before (Foldi et al., 2017; McIlroy et al., 2013). To pull down Wek-FLAG, S2 cells were lysed in ice-cold NP-40 lysis buffer (50mM Tris-HCl pH:8.0, 150mM NaCl, 1% Igepal-630) supplemented with protease inhibitor cocktail (Pierce). 10% of the lysate was aliquoted and used as input. Immunoprecipitation of expressed Wek-Flag was carried out using anti-FLAG^R^ M2 Magnetic Beads (Sigma-Aldrich/Merck, Cat. No. M8823) according to manufacturers’ instructions. The bait-loaded beads were incubated with fresh lysate from untransfected S2 cells for 1 h at 4 °C to capture potential endogenous interactors. After washes, bound and unbound fractions were analyzed by SDS-PAGE and Western blot. Beads-only and FLAG peptide blocking controls were included. Proteins were visualised on SDS-Page gels by Coomassie Blue staining, following by washing in ddH20, and each gel band was cut into several pieces; alternatively, the protein bands of interest were excised. Gel fragments were analyzed by liquid chromatography–tandem mass spectrometry (LC–MS/MS) at the Proteomics Facility at the University of Birmingham. Analysis of hits and identification of GO terms was carried out using Metascape (https://metascape.org/gp/index.html#/main/step1) (Zhou et al., 2019).

#### Phostag gel from S2 cells

To analyse phosphorylation profiles of Wek, wild-type and mutant forms of *wek* - *pAct-Wek-FLAG, pAct-WekG2A-FLAG, pAct-WekS6N-FLAG, pAct-WekS164A-FLAG* or *pAct-WekS168A-FLAG* - were expressed in S2 cells as described above. To detect phosphorylated Wek and Wek mutants in SDS-PAGE, we used Phos-tag AAL-107 (FMS Laboratory) according to the manufactureŕs instructions. The phostag gels, using Mn^2+^-Phos-tag SDS-PAGE, were prepared as described by Kinoshita (Kinoshita et al., 2009). Optimization included Phos-tag concentration (typically 20-30 uM), and MnCl2 (60 uM), acrylamide percentage 10 % and electrophoresis run for 1.15h. This was followed by EDTA wash, membrane transfer and western blotting, as described below.

#### Co-immuno-precipitation

S2 cells were transfected with wild-type *pAct-wek-FLAG*, as described above. Following a 48-hour incubation period, cells were harvested and washed in PBS. Cell pellets were lysed in 600μL NP-40 buffer (50mM Tris-HCl pH 8.0, 150mM NaCl, 1% Igepal-630) supplemented with protease inhibitor cocktail (Pierce). Lysates were incubated with anti-Flag antibody-conjugated magnetic beads (Sigma-Aldrich) at room temperature for 2 hours. Thereafter, the beads were thoroughly washed in Tris-buffered saline with Tween-20 buffer. The proteins were then eluted in 40μL of 2x Laemmli buffer, run on an SDS-PAGE gel and analysed by Western blot, as described below. Anti-Yki was used to detect Yki, from naturally and endogenously expressed *yki* in S2 cells, in western blots (see below).

#### Western blotting

Tagged Wek protein samples were run in SDS-PAGE, and following membrane transfer, Western blotting was carried out according to standard procedures. Primary antibodies used were rabbit anti-Flag (1:2000; F7425, Sigma Aldrich), chicken anti-HA (12CA2; 1:2000; 11 583 816 001; Roche), mouse anti-V5 (1:5000; Invitrogen; #R960-25) and, rabbit anti-Yki (1:400 gift from Irvine lab). Secondary antibodies used were anti-rabbit HRP (1:5000; PI-1000; Vector Laboratories), anti-chicken HRP (1:10000; 703-035-155; Jackson ImmnunoResearch Laboratories, Inc) and anti-mouse HRP (1:5000; PI-2000 Vector Laboratories). After incubation with primary and secondary antibodies, protein bands were visualized using SuperSignal™ Western Blot Chemiluminescent Substrate (Thermo Fisher Scientific num. cat. 34579) following manufacturer’s protocol.

#### Cell biology Immunostaining

To visualise Wek intracellular trafficking in S2 cells, we transfected them with wild-type and mutant forms of *wek* - *pAct-Wek-FLAG, pAct-Wek^G2A^, FLAG, pAct-Wek^S6N^-Flag, pAct-Wek^S164A^-Flag* or *pAct-Wek^S168A^-Flag*, as described above. *Drosophila* Scheneider 2 (S2) cells were cultured under standard conditions and seeded onto poly-L-lysine-coated coverslips to promote adherence. Cells were fixed with 4% paraformaldehyde in phosphate-buffered saline (PBS) for 10 minutes at room temperature. Following fixation, cells were permeabilized with 0.1% Triton X-100 in PBS for 5 minutes and blocked with 5% milk in PBS for 1 hour at room temperature to reduce nonspecific binding, followed by permeabilization and blocking using established procedures. Primary antibodies used were: mouse monoclonal anti-FLAG M2 (1:2000; F1804; Sigma Aldrich); secondary antibody: FITC-conjugated anti-Mouse at 1:1000 (Thermo Fisher Scientific). Nuclei were counterstained with DAPI and mounted in antifade Vectashield medium (Vector Laboratories, Cat. No. H-1200).

For stainings in larval fat body, flies were reared constantly at 25°C from egg-laying until L3 wandering, unless otherwise indicated. Fat bodies were dissected from 120h AEL old larvae, in ice-cold PBS and fixed in 4% PFA at room temperature for 20 minutes, then washed with 0.3% PBT-Triton for 4 times followed by standard staining procedures. Stained tissues were mounted in Vectashield containing DAPI on poly-lysine coated slides. For stainings in pupal and adult brains, flies were reared constantly at 25°C until 8h or 24h after puparium formation (APF) or 48h after eclosion, and brains were dissected in ice-cold PBS and fixed in 2% PFA at room temperature for 40 minutes. Fixed brains were then washed in 0.3% PBT-Triton for pupal brains or 0.5% PBT-Triton for 5 times. Antibody staining was carried out following standard procedures. Primary antibodies used were: rabbit-anti-GFP at 1:250 (Molecular Probes); Rat anti-HA at 1:100 (Roche, Merck); Mouse anti-V5 at 1:100 (Thermo Fisher); Rat anti-Flag at 1:100 (Novus Biologicals); anti-pH3 at 1:250 (Upstate Biotechnology); Rabbit anti-Yki (gift from KD Irvine); Secondary antibodies used were: Alexa Donkey-anti-Rabbit 488 at 1:250; Alexa Goat-anti-Mouse 488 at 1:250; Alexa Goat-anti-Mouse 647 at 1:250; Alexa Goat-anti-Rat 488 at 1:250.

### Microscopy and Imaging

#### Fluorescence laser scanning confocal microscopy

Fluorescence microscopy images of S2 cells were acquired on Zeiss LSM 710 confocal microscope with a 25x oil immersion lens, at a resolution of 1024 × 1024 pixels. Negative controls included omission of the primary antibody and staining of un-transfected cells. Fat body cells were imaged using a Zeiss LSM900 or Leica Stellaris laser scanning confocal microscopes, 63x/1.4 oil lens was used for larval fat body, with step size 1µm, and resolution of 1024 x 1024. Imaging of pupal and adult fly brains was carried out using a Leica SP8 laser scanning confocal microscope, 20x oil lens was used for whole adult brains or pupal brains at zoom 1.0; 40x oil lens was used for antennal lobes at zoom 0.9; with step size: whole brain 0.96µm, antennal lobes 0.5µm; acquisition speed was 400Hz, air unit 1, resolution of 1024 x 1024, and no line averaging was applied. For non-quantitative data, confocal stacks of optical sections were processed using ImageJ/Fiji software, and figures were created using Adobe Photoshop and Illustrator.

#### Automatic cell counting with DeadEasy

For automatic counting of Histone-YFP labelled cells in the adult central brain, the DeadEasy Central Brain ImageJ plug-in (Li et al., 2020) was used. Raw stacks of confocal images were used, and a Region of Interest (ROI) was selected for the cell counting throughout the 3D-stack. All the parameters were maintained the same as published with no adjustment.

#### Cell counting with Imaris

For the supplementary data Figure S4, YFP and anti-phospho-Histone-H3 (pH3) cells in pupal central brain were counted using Imaris. “Surface” module was used to define the ROI of central brain and the same ROI was applied for both YFP and PH3 counting for each brain. The ROI was masked to make sure the cell counting was restricted within the central brain. “Spots” module was used for cell counting. In 8h pupal brain, “Estimated XY Diameter” was set as 5.68µm and “Background Subtraction” was selected. “Classify Spots” filter was set as “Quality above 22.0” for PH3 cells and “Quality above 15.0” for YFP cells. In 24h pupal brain, “Estimated XY Diameter” was set as 8µm for PH3 cells and 5µm for YFP cells. “Classify Spots” filter was set as “Quality above 27.0” for PH3 cells and “Quality above 15.0” for YFP cells. The “Edit” function was used to correct any false counting detected. The “Total Number of Spots” were collected to compare.

#### Volume analysis with Imaris

To analyse glial cell membrane volume in adult antennal lobes, laser scanning confocal microscopy image data were processed with Imaris using the “Surface” module. An ROI was drawn in 3D, by manually tracing the outline of each antennal lobe, throughout the confocal optical sections. The ROI was then masked to make sure the analysis was just applied within the ROI. For the segmentation, “Smooth” was selected and was set as 0.710µm; “Background Subtraction” was selected and as set as 2.66µm; the “Threshold” for Background Subtraction was set automatically by the software (and could vary across samples due to the variability of immunostaining and image acquisition); no filter for “Classify Surfaces” was selected. The “Edit” function was used to correct any inaccuracies detected. The “SUM” value of signal volume was collected and compared across genotypes. The “signal volume” corresponds to the segmented glial expression of mCD8GFP in the antennal lobes, in 3D.

#### Statistical analysis

For statistical analyses, we used GraphPad Prism, with significance at p<0.05 (confidence interval 95%). Continuous numerical data were tested for normality (e.g. Kolmogorov-Smirnov test) and equality of variance (e.g. Levene’s test). If data were normally distributed, a Student t-test was used when comparing two groups, whereas a One Way ANOVA test was used when comparing more than two groups, which was followed by Tukey’s tests for multiple comparisons corrections to compare all groups to each other.

## AUTHOR CONTRIBUTIONS

MDPS, GL, MMon, FRC, MyMa, EC, MMor executed experiments and curated data; MDPS, GL, MMon, FRC, NJG and AH designed experiments and analysed data; AP and RF carried out analysis; AH, NJG, RF supervised; AH and NJG secured funding for the project; AH conceived and directed the project; AH and MDPS wrote the manuscript; all authors contributed to improving the manuscript.

## ACKNOWLEDGEMENTS

We thank all the members of our the Hidalgo Lab for comments on the manuscript; Peter Kean for bioinformatics advice; Kieran Harvey, Ken Irvine, Nic Tapon and Barry Thompson for kind gifts of antibodies and flies; Developmental Studies Hybridoma Bank, for antibodies; AddGene for plasmids; FlyBase for unlimited assistance. This project was funded by BBSRC Project Grant BB/R00871X/1 to AH and NJG, Wellcome Trust Investigator Award to AH and BBSRC MIBTP PhD studentships to AP, EC and ES.

---

**Supplementary Figure S1 Dynamic Light Scattering detailed evidence that Wek can directly interact with Toll-2 and Toll-6.** Dynamic Light Scattering (DLS) analysis of particle size distributions under different conditions. Representative correlograms g(τ) (top), residuals of the fit (middle), and intensity-weighted size distributions (bottom) are shown for samples from Wek (11–13), Wek + Toll2 (17–19), and Wek + T6 (14–16). The hydrodynamic radius (Rh) distributions indicate that Wek displays an increased Rh in the presence of both Toll2 and Toll6, reflecting changes in particle populations depending on the experimental condition.

**Supplementary Figure S2 Bulk RNAseq upon wek over-expression: RNAseq Go term and KEGG term analyses (A)** Over-expression of *wek-HA* in MyD88+ cells in adult heads (*tubGAL980^ts^, MyD88> wek* vs. *tubGAL980^ts^, MyD88>+ control*) resulted in both up- and down-regulation of gene expression. KEGG and GO term analysis showed that Toll signalling and innate immunity related genes were dramatically downregulated. **(B)** Over-expression of *wek-HA* together with *dilp-6* in glial cells *(tubGAL980^ts^, repo>dilp-6, wek* vs. *tubGAL980^ts^, repo>dilp-6* control) also up- and down-regulated gene expression. Up-regulated genes included those involved in oxidation-reduction, response to stress, lipid metabolism, cell proliferation, ribosome biogenesis and translation.

**Supplementary Figure S3 Targets of Wek identified with ChipSeq. (A)** GO term analysis of significant Wek targets identified with ChipSeq, p-values in **(B),** shown in Metascape representation. No antimicrobial genes were found amongst target genes, which instead included genes involved in stress response, autophagy, cell division, stem cells, metabolism, RNA biology and neuronal functions.

**Supplementary Figure S4 Wek promotes ensheathing glia proliferation during pupal development. (A,C)** Co-localisation of the mitotic marker anti-phospho-Histone-H3 and Histone-HYP in NP6520Gal4+ glial cells, at 8h after puparium formation (APF). Cells over-expressing *wek-HA* can be larger than controls, have pH3, but cell number has not yet increased as cells are still growing and dividing. **(B,D)** by 24h APF, cell division lead to an increase in cell number in brain over-expressing *wek-HA* compared to wild-type controls *(NP6520>his-YFP, wek-HA).* Mann Whitney U test, *p<0.05. See Table S11 for genotype details and sample sizes.

**Supplementary Figure S5 Annotation of human proteins containing a ZAD domain identifies products of the single *ZFP276* gene.** Interpro output of human ZAD-containing gene products.

**Table S1 S2 cell pull-down Wek-FLAG with anti-FLAG.** List of proteins identified through binding of isolated Wek-FLAG to S2-cell lysate proteins. Identification of Sarm as a Wek-interacting protein serves as a positive control. Insulin signalling factors and Yki are amongst the pull-down hits.

**Table S2 RNAseq down-regulated genes from MyD88 cells over-expressing *wek-HA*.** Bulk RNAseq list of significantly down-regulated genes in heads of flies over-expressing *wek* in MyD88+ cells (ie *tubGAL80ts, MyD88>wek-HA vs. tubGAL80ts, MyD88>+* controls) restricted to the adult.

**Table S3 RNAseq up-regulated genes from MyD88 cells over-expressing *wek-HA*.** Bulk RNAseq list of significantly up-regulated genes in heads of flies over-expressing *wek* in MyD88+ cells (ie *tubGAL80ts, MyD88>wek-HA vs. tubGAL80ts, MyD88>+* controls) restricted to the adult.

**Table S4 RNAseq down-regulated genes from glial cells over-expressing *dilp-6* and *wek-HA*.** Bulk RNAseq list of significantly down-regulated genes in heads of flies over-expressing *wek-HA* in glial cells (ie *tubGAL80ts, repo>dilp-6, wek-HA vs. tubGAL80ts, repo>dilp-6* controls) restricted to the adult.

**Table S5 RNAseq up-regulated genes from glial cells over-expressing *dilp-6* and *wek-HA*.** Bulk RNAseq list of significantly up-regulated genes in heads of flies over-expressing *wek* in glial cells (ie *tubGAL80ts, repo>dilp-6, wek-HA vs. tubGAL80ts, repo>dilp-6* controls) restricted to the adult.

**Table S6 ChIPseq wek targets.** Wek target genes identified across four biological replicates from dissected adult brains, and two biological replicates from dissected larval central nervous system.

**Table S7 Common Wek targets in RNAseq and Chiseq.**

**Table S8 Wek DNA-binding motifs within target genes identified with ChipSeq**

**Table S9 Similarity of ChipSeq Wek binding sites to those of other transcription factors.**

**Table S10 Genotypes and sample sizes for in vivo imaging data.** Full genotypes and sample sizes for genetic analysis with imaging samples in Figures 3F,G, 4C-F, 6A,B and 7A-F.

**Table S11 List of *Drosophila* stocks used or generated.**

**Table S12 List of constructs and primers used for molecular cloning. Table S13 Putative *wek* orthologues identified with PhylomeDB.**

## REFERENCES

Abarca-Merlin, D.M., Martinez-Duran, J.A., Medina-Perez, J.D., Rodriguez-Santos, G., and Alvarez-Arellano, L. (2024). From Immunity to Neurogenesis: Toll-like Receptors as Versatile Regulators in the Nervous System. Int J Mol Sci 25.

Aberle, T., Piefke, S., Hillgartner, S., Tamm, E.R., Wegner, M., and Kuspert, M. (2022). Transcription factor Zfp276 drives oligodendroglial differentiation and myelination by switching off the progenitor cell program. Nucleic Acids Res 50, 1951–1968.

Alonso, R., Pisa, D., Marina, A.I., Morato, E., Rabano, A., and Carrasco, L. (2014). Fungal infection in patients with Alzheimer’s disease. J Alzheimers Dis 41, 301–311.

Angileri, K.M., Bagia, N.A., and Feschotte, C. (2022). Transposon control as a checkpoint for tissue regeneration. Development 149.

Anthoney, N., Foldi, I., and Hidalgo, A. (2018). Toll and Toll-like receptor signalling in development. Development 145.

Belinda, L.W., Wei, W.X., Hanh, B.T., Lei, L.X., Bow, H., and Ling, D.J. (2008). SARM: a novel Toll-like receptor adaptor, is functionally conserved from arthropod to human. Mol Immunol 45, 1732–1742.

Birey, F., Kokkosis, A.G., and Aguirre, A. (2017). Oligodendroglia-lineage cells in brain plasticity, homeostasis and psychiatric disorders. Curr Opin Neurobiol 47, 93–103.

Bock, K. (2021). Brain Inflamed: Uncovering the hidden causes of anxiety, depression and other mood disorders in adolescents and teens (Piatkus).

Bonchuk, A., Boyko, K., Fedotova, A., Nikolaeva, A., Lushchekina, S., Khrustaleva, A., Popov, V., and Georgiev, P. (2021). Structural basis of diversity and homodimerization specificity of zinc-finger-associated domains in Drosophila. Nucleic Acids Res 49, 2375–2389.

Carty, M., and Bowie, A.G. (2019). SARM: From immune regulator to cell executioner. Biochem Pharmacol 161, 52–62.

Carty, M., Goodbody, R., Schroder, M., Stack, J., Moynagh, P.N., and Bowie, A.G. (2006). The human adaptor SARM negatively regulates adaptor protein TRIF-dependent Toll-like receptor signaling. Nat Immunol 7, 1074–1081.

Chauhan, M., Martinak, P.E., Hollenberg, B.M., and Goodman, A.G. (2024). Drosophila melanogaster Toll-9 elicits antiviral immunity against Drosophila C virus. bioRxiv.

Chen, C.Y., Shih, Y.C., Hung, Y.F., and Hsueh, Y.P. (2019). Beyond defense: regulation of neuronal morphogenesis and brain functions via Toll-like receptors. J Biomed Sci 26, 90.

Chen, L.Y., Wang, J.C., Hyvert, Y., Lin, H.P., Perrimon, N., Imler, J.L., and Hsu, J.C. (2006). Weckle is a zinc finger adaptor of the toll pathway in dorsoventral patterning of the Drosophila embryo. Curr Biol 16, 1183–1193.

Chung, H.R., Lohr, U., and Jackle, H. (2007). Lineage-specific expansion of the zinc finger associated domain ZAD. Mol Biol Evol 24, 1934–1943.

Chung, H.R., Schafer, U., Jackle, H., and Bohm, S. (2002). Genomic expansion and clustering of ZAD-containing C2H2 zinc-finger genes in Drosophila. EMBO Rep 3, 1158–1162.

Church, J.S., Kigerl, K.A., Lerch, J.K., Popovich, P.G., and McTigue, D.M. (2016). TLR4 Deficiency Impairs Oligodendrocyte Formation in the Injured Spinal Cord. J Neurosci 36, 6352–6364.

Collins, T., Stone, J.R., and Williams, A.J. (2001). All in the family: the BTB/POZ, KRAB, and SCAN domains. Mol Cell Biol 21, 3609–3615.

Davis, J.R., and Tapon, N. (2019). Hippo signalling during development. Development 146.

Ding, R., Weynans, K., Bossing, T., Barros, C.S., and Berger, C. (2016). The Hippo signalling pathway maintains quiescence in Drosophila neural stem cells. Nat Commun 7, 10510.

Ding, W.Y., Huang, J., and Wang, H. (2020). Waking up quiescent neural stem cells: Molecular mechanisms and implications in neurodevelopmental disorders. PLoS Genet 16, e1008653.

Donnelly, C.R., Chen, O., and Ji, R.R. (2020). How Do Sensory Neurons Sense Danger Signals? Trends Neurosci 43, 822–838.

Ecco, G., Imbeault, M., and Trono, D. (2017). KRAB zinc finger proteins. Development 144, 2719–2729.

Edelstein, L.C., and Collins, T. (2005). The SCAN domain family of zinc finger transcription factors. Gene 359, 1–17.

Essuman, K., Summers, D.W., Sasaki, Y., Mao, X., DiAntonio, A., and Milbrandt, J. (2017). The SARM1 Toll/Interleukin-1 Receptor Domain Possesses Intrinsic NAD(+) Cleavage Activity that Promotes Pathological Axonal Degeneration. Neuron 93, 1334–1343 e1335.

Fan, W., Jurado-Arjona, J., Alanis-Lobato, G., Peron, S., Berger, C., Andrade-Navarro, M.A., Falk, S., and Berninger, B. (2023). The transcriptional co-activator Yap1 promotes adult hippocampal neural stem cell activation. EMBO J 42, e110384.

Feldman, D.E., and Brecht, M. (2005). Map plasticity in somatosensory cortex. Science 310, 810–815.

Foldi, I., Anthoney, N., Harrison, N., Gangloff, M., Verstak, B., Nallasivan, M.P., AlAhmed, S., Zhu, B., Phizacklea, M., Losada-Perez, M., et al. (2017). Three-tier regulation of cell number plasticity by neurotrophins and Tolls in Drosophila. J Cell Biol 216, 1421–1438.

Gage, F.H. (2019). Adult neurogenesis in mammals. Science 364, 827–828.

Gailite, I., Aerne, B.L., and Tapon, N. (2015). Differential control of Yorkie activity by LKB1/AMPK and the Hippo/Warts cascade in the central nervous system. Proc Natl Acad Sci U S A 112, E5169–5178.

Gay, N.J., and Gangloff, M. (2007). Structure and function of Toll receptors and their ligands. Annu Rev Biochem 76, 141–165.

Gilley, J., Jackson, O., Pipis, M., Estiar, M.A., Al-Chalabi, A., Danzi, M.C., van Eijk, K.R., Goutman, S.A., Harms, M.B., Houlden, H., et al. (2021). Enrichment of SARM1 alleles encoding variants with constitutively hyperactive NADase in patients with ALS and other motor nerve disorders. Elife 10.

Goulev, Y., Fauny, J.D., Gonzalez-Marti, B., Flagiello, D., Silber, J., and Zider, A. (2008). SCALLOPED interacts with YORKIE, the nuclear effector of the hippo tumor-suppressor pathway in Drosophila. Curr Biol 18, 435–441.

Grand, R.J. (1989). Acylation of viral and eukaryotic proteins. Biochem J 258, 625–638.

Hoffmann, J.A. (2003). The immune response of Drosophila. Nature 426, 33–38.

Hoffmann, J.A., and Reichhart, J.M. (2002). Drosophila innate immunity: an evolutionary perspective. Nat Immunol 3, 121–126.

Holtmaat, A., and Svoboda, K. (2009). Experience-dependent structural synaptic plasticity in the mammalian brain. Nat Rev Neurosci 10, 647–658.

Huang, J., Wu, S., Barrera, J., Matthews, K., and Pan, D. (2005). The Hippo signaling pathway coordinately regulates cell proliferation and apoptosis by inactivating Yorkie, the Drosophila Homolog of YAP. Cell 122, 421–434.

Hung, Y.F., Chen, C.Y., Shih, Y.C., Liu, H.Y., Huang, C.M., and Hsueh, Y.P. (2018). Endosomal TLR3, TLR7, and TLR8 control neuronal morphology through different transcriptional programs. J Cell Biol 217, 2727–2742.

Imler, J.L., and Hoffmann, J.A. (2001). Toll receptors in innate immunity. Trends Cell Biol 11, 304–311.

Jauch, R., Bourenkov, G.P., Chung, H.R., Urlaub, H., Reidt, U., Jackle, H., and Wahl, M.C. (2003). The zinc finger-associated domain of the Drosophila transcription factor grauzone is a novel zinc-coordinating protein-protein interaction module. Structure 11, 1393–1402.

Jumper, J., Evans, R., Pritzel, A., Green, T., Figurnov, M., Ronneberger, O., Tunyasuvunakool, K., Bates, R., Zidek, A., Potapenko, A., et al. (2021). Highly accurate protein structure prediction with AlphaFold. Nature 596, 583–589.

Karnik, A., and Joshi, A. (2025). SARM1: The Checkpoint of Axonal Degeneration in the Nervous System Disorders. Mol Neurobiol 62, 9240–9257.

Kato, K., Orihara-Ono, M., and Awasaki, T. (2020). Multiple lineages enable robust development of the neuropil-glia architecture in adult Drosophila. Development 147.

Kinoshita, E., Kinoshita-Kikuta, E., and Koike, T. (2009). Separation and detection of large phosphoproteins using Phos-tag SDS-PAGE. Nat Protoc 4, 1513–1521.

Krystel, J., and Ayyanathan, K. (2013). Global analysis of target genes of 21 members of the ZAD transcription factor family in Drosophila melanogaster. Gene 512, 373–382.

Kumar, V. (2019). Toll-like receptors in the pathogenesis of neuroinflammation. J Neuroimmunol 332, 16–30.

Kwon, Y., Song, W., Droujinine, I.A., Hu, Y., Asara, J.M., and Perrimon, N. (2015). Systemic organ wasting induced by localized expression of the secreted insulin/IGF antagonist ImpL2. Dev Cell 33, 36–46.

Laue, T.M., Shah, B.D., Ridgeway, T.M., and Pelletier, S.L. (1992). Computer-aided interpretation of analytical sedimentation data for proteins. In Analytical Ultracentrifugation in Biochemistry and Polymer Science, e. S.E. Harding and J.C. Horton, ed. (Royal Society of Chemistry)), pp. 90–125.

Lauenstein, J.U., Udgata, A., Bartram, A., De Sutter, D., Fisher, D.I., Halabi, S., Eyckerman, S., and Gay, N.J. (2019). Phosphorylation of the multifunctional signal transducer B-cell adaptor protein (BCAP) promotes recruitment of multiple SH2/SH3 proteins including GRB2. J Biol Chem 294, 19852–19861.

Lemaitre, B., Nicolas, E., Michaut, L., Reichhart, J.M., and Hoffmann, J.A. (1996). The dorsoventral regulatory gene cassette spatzle/Toll/cactus controls the potent antifungal response in Drosophila adults. Cell 86, 973–983.

Leulier, F., and Lemaitre, B. (2008). Toll-like receptors--taking an evolutionary approach. Nat Rev Genet 9, 165–178.

Li, G., Forero, M.G., Wentzell, J.S., Durmus, I., Wolf, R., Anthoney, N.C., Parker, M., Jiang, R., Hasenauer, J., Strausfeld, N.J., et al. (2020). A Toll-receptor map underlies structural brain plasticity. Elife 9.

Li, G., and Hidalgo, A. (2021). The Toll Route to Structural Brain Plasticity. Front Physiol 12, 679766.

Li, Y., Chen, L., Zhao, W., Sun, L., Zhang, R., Zhu, S., Xie, K., Feng, X., Wu, X., Sun, Z., et al. (2021). Food reward depends on TLR4 activation in dopaminergic neurons. Pharmacol Res 169, 105659.

Lin, C.W., and Hsueh, Y.P. (2014). Sarm1, a neuronal inflammatory regulator, controls social interaction, associative memory and cognitive flexibility in mice. Brain Behav Immun 37, 142–151.

Lin, C.W., Liu, H.Y., Chen, C.Y., and Hsueh, Y.P. (2014). Neuronally-expressed Sarm1 regulates expression of inflammatory and antiviral cytokines in brains. Innate Immun 20, 161–172.

Liu, B., Zheng, Y., Yin, F., Yu, J., Silverman, N., and Pan, D. (2016). Toll Receptor-Mediated Hippo Signaling Controls Innate Immunity in Drosophila. Cell 164, 406–419.

Lu, B., Pang, P.T., and Woo, N.H. (2005). The yin and yang of neurotrophin action. Nat Rev Neurosci 6, 603–614.

Manning, S.A., Kroeger, B., and Harvey, K.F. (2020). The regulation of Yorkie, YAP and TAZ: new insights into the Hippo pathway. Development 147.

McIlroy, G., Foldi, I., Aurikko, J., Wentzell, J.S., Lim, M.A., Fenton, J.C., Gay, N.J., and Hidalgo, A. (2013). Toll-6 and Toll-7 function as neurotrophin receptors in the Drosophila melanogaster CNS. Nat Neurosci 16, 1248–1256.

McLaughlin, C.N., Nechipurenko, I.V., Liu, N., and Broihier, H.T. (2016). A Toll receptor-FoxO pathway represses Pavarotti/MKLP1 to promote microtubule dynamics in motoneurons. J Cell Biol 214, 459–474.

McLaughlin, C.N., Perry-Richardson, J.J., Coutinho-Budd, J.C., and Broihier, H.T. (2019). Dying Neurons Utilize Innate Immune Signaling to Prime Glia for Phagocytosis during Development. Dev Cell 48, 506–522 e506.

Mirdita, M., Schutze, K., Moriwaki, Y., Heo, L., Ovchinnikov, S., and Steinegger, M. (2022). ColabFold: making protein folding accessible to all. Nat Methods 19, 679–682.

Mo, J.S., Park, H.W., and Guan, K.L. (2014). The Hippo signaling pathway in stem cell biology and cancer. EMBO Rep 15, 642–656.

Mukherjee, P., Winkler, C.W., Taylor, K.G., Woods, T.A., Nair, V., Khan, B.A., and Peterson, K.E. (2015). SARM1, Not MyD88, Mediates TLR7/TLR9-Induced Apoptosis in Neurons. J Immunol 195, 4913-4921.

Nakamoto, M., Moy, R.H., Xu, J., Bambina, S., Yasunaga, A., Shelly, S.S., Gold, B., and Cherry, S. (2012). Virus recognition by Toll-7 activates antiviral autophagy in Drosophila. Immunity 36, 658–667.

Narbonne-Reveau, K., Charroux, B., and Royet, J. (2011). Lack of an antibacterial response defect in Drosophila Toll-9 mutant. PLoS One 6, e17470.

Oh, H., and Irvine, K.D. (2008). In vivo regulation of Yorkie phosphorylation and localization. Development 135, 1081–1088.

Oh, H., Slattery, M., Ma, L., Crofts, A., White, K.P., Mann, R.S., and Irvine, K.D. (2013). Genome-wide association of Yorkie with chromatin and chromatin-remodeling complexes. Cell Rep 3, 309–318.

Okada, T., Maeda, A., Iwamatsu, A., Gotoh, K., and Kurosaki, T. (2000). BCAP: the tyrosine kinase substrate that connects B cell receptor to phosphoinositide 3-kinase activation. Immunity 13, 817–827.

Okun, E., Griffioen, K., Barak, B., Roberts, N.J., Castro, K., Pita, M.A., Cheng, A., Mughal, M.R., Wan, R., Ashery, U., et al. (2010). Toll-like receptor 3 inhibits memory retention and constrains adult hippocampal neurogenesis. Proc Natl Acad Sci U S A 107, 15625–15630.

Okun, E., Griffioen, K.J., Lathia, J.D., Tang, S.C., Mattson, M.P., and Arumugam, T.V. (2009). Toll-like receptors in neurodegeneration. Brain Res Rev 59, 278–292.

Okun, E., Griffioen, K.J., and Mattson, M.P. (2011). Toll-like receptor signaling in neural plasticity and disease. Trends Neurosci 34, 269–281.

Osterloh, J.M., Yang, J., Rooney, T.M., Fox, A.N., Adalbert, R., Powell, E.H., Sheehan, A.E., Avery, M.A., Hackett, R., Logan, M.A., et al. (2012). dSarm/Sarm1 is required for activation of an injury-induced axon death pathway. Science 337, 481–484.

Pare, A.C., Vichas, A., Fincher, C.T., Mirman, Z., Farrell, D.L., Mainieri, A., and Zallen, J.A. (2014). A positional Toll receptor code directs convergent extension in Drosophila. Nature 515, 523–527.

Park, H., and Poo, M.M. (2013). Neurotrophin regulation of neural circuit development and function. Nat Rev Neurosci 14, 7–23.

Park, S.J., Lee, J.Y., Kim, S.J., Choi, S.Y., Yune, T.Y., and Ryu, J.H. (2015). Toll-like receptor-2 deficiency induces schizophrenia-like behaviors in mice. Sci Rep 5, 8502.

Parker, J., and Struhl, G. (2015). Scaling the Drosophila Wing: TOR-Dependent Target Gene Access by the Hippo Pathway Transducer Yorkie. PLoS Biol 13, e1002274.

Pascual, M., Calvo-Rodriguez, M., Nunez, L., Villalobos, C., Urena, J., and Guerri, C. (2021). Toll-like receptors in neuroinflammation, neurodegeneration, and alcohol-induced brain damage. IUBMB Life 73, 900–915.

Peng, J., Yuan, Q., Lin, B., Panneerselvam, P., Wang, X., Luan, X.L., Lim, S.K., Leung, B.P., Ho, B., and Ding, J.L. (2010). SARM inhibits both TRIF- and MyD88-mediated AP-1 activation. Eur J Immunol 40, 1738–1747.

Pisa, D., Alonso, R., Juarranz, A., Rabano, A., and Carrasco, L. (2015). Direct visualization of fungal infection in brains from patients with Alzheimer’s disease. J Alzheimers Dis 43, 613–624.

Poon, C.L., Mitchell, K.A., Kondo, S., Cheng, L.Y., and Harvey, K.F. (2016). The Hippo Pathway Regulates Neuroblasts and Brain Size in Drosophila melanogaster. Curr Biol 26, 1034–1042.

Reddy, B.V., and Irvine, K.D. (2011). Regulation of Drosophila glial cell proliferation by Merlin-Hippo signaling. Development 138, 5201–5212.

Rolls, A., Shechter, R., London, A., Ziv, Y., Ronen, A., Levy, R., and Schwartz, M. (2007). Toll-like receptors modulate adult hippocampal neurogenesis. Nat Cell Biol 9, 1081–1088.

Roth, S.W., Bitterman, M.D., Birnbaum, M.J., and Bland, M.L. (2018). Innate Immune Signaling in Drosophila Blocks Insulin Signaling by Uncoupling PI(3,4,5)P(3) Production and Akt Activation. Cell Rep 22, 2550–2556.

Rowe, D.C., McGettrick, A.F., Latz, E., Monks, B.G., Gay, N.J., Yamamoto, M., Akira, S., O’Neill, L.A., Fitzgerald, K.A., and Golenbock, D.T. (2006). The myristoylation of TRIF-related adaptor molecule is essential for Toll-like receptor 4 signal transduction. Proc Natl Acad Sci U S A 103, 6299–6304.

Sakakibara, Y., Yamashiro, R., Chikamatsu, S., Hirota, Y., Tsubokawa, Y., Nishijima, R., Takei, K., Sekiya, M., and Iijima, K.M. (2023). Drosophila Toll-9 is induced by aging and neurodegeneration to modulate stress signaling and its deficiency exacerbates tau-mediated neurodegeneration. iScience 26, 105968.

Sanchez-Petidier, M., Guerri, C., and Moreno-Manzano, V. (2022). Toll-like receptors 2 and 4 differentially regulate the self-renewal and differentiation of spinal cord neural precursor cells. Stem Cell Res Ther 13, 117.

Schuck, P. (2003). On the analysis of protein self-association by sedimentation velocity analytical ultracentrifugation. Anal Biochem 320, 104–124.

Seelig, S., Ryus, C.R., Harrison, R.F., Wilson, M.P., and Wong, A.H. (2019). Cryptococcal Meningoencephalitis Presenting as a Psychiatric Emergency. J Emerg Med 57, 203–206.

Shechter, R., Ronen, A., Rolls, A., London, A., Bakalash, S., Young, M.J., and Schwartz, M. (2008). Toll-like receptor 4 restricts retinal progenitor cell proliferation. J Cell Biol 183, 393–400.

Shen, Y., Qin, H., Chen, J., Mou, L., He, Y., Yan, Y., Zhou, H., Lv, Y., Chen, Z., Wang, J., et al. (2016). Postnatal activation of TLR4 in astrocytes promotes excitatory synaptogenesis in hippocampal neurons. J Cell Biol 215, 719–734.

Shi, F.D., and Yong, V.W. (2025). Neuroinflammation across neurological diseases. Science 388, eadx0043.

Singh, D.N.D., Roberts, A.R.E., Wang, X., Li, G., Quesada Moraga, E., Alliband, D., Ballou, E., Tsai, H.J., and Hidalgo, A. (2025). Toll-1-dependent immune evasion induced by fungal infection leads to cell loss in the Drosophila brain. PLoS Biol 23, e3003020.

Squillace, S., and Salvemini, D. (2022). Toll-like receptor-mediated neuroinflammation: relevance for cognitive dysfunctions. Trends Pharmacol Sci 43, 726–739.

Stetefeld, J., McKenna, S.A., and Patel, T.R. (2016). Dynamic light scattering: a practical guide and applications in biomedical sciences. Biophys Rev 8, 409–427.

Strassburger, K., Tiebe, M., Pinna, F., Breuhahn, K., and Teleman, A.A. (2012). Insulin/IGF signaling drives cell proliferation in part via Yorkie/YAP. Dev Biol 367, 187–196.

Sun, J., Rojo-Cortes, F., Ulian-Benitez, S., Forero, M.G., Li, G., Singh, D.N.D., Wang, X., Cachero, S., Moreira, M., Kavanagh, D., et al. (2024). A neurotrophin functioning with a Toll regulates structural plasticity in a dopaminergic circuit. Elife 13.

Suzawa, M., Muhammad, N.M., Joseph, B.S., and Bland, M.L. (2019). The Toll Signaling Pathway Targets the Insulin-like Peptide Dilp6 to Inhibit Growth in Drosophila. Cell Rep 28, 1439–1446 e1435.

Tamada, M., Shi, J., Bourdot, K.S., Supriyatno, S., Palmquist, K.H., Gutierrez-Ruiz, O.L., and Zallen, J.A. (2021). Toll receptors remodel epithelia by directing planar-polarized Src and PI3K activity. Dev Cell 56, 1589–1602 e1589.

Tauszig, S., Jouanguy, E., Hoffmann, J.A., and Imler, J.L. (2000). Toll-related receptors and the control of antimicrobial peptide expression in Drosophila. Proc Natl Acad Sci U S A 97, 10520–10525.

Thelen, M., Rosen, A., Nairn, A.C., and Aderem, A. (1991). Regulation by phosphorylation of reversible association of a myristoylated protein kinase C substrate with the plasma membrane. Nature 351, 320–322.

Towler, D.A., Gordon, J.I., Adams, S.P., and Glaser, L. (1988). The biology and enzymology of eukaryotic protein acylation. Annu Rev Biochem 57, 69–99.

Troubat, R., Barone, P., Leman, S., Desmidt, T., Cressant, A., Atanasova, B., Brizard, B., El Hage, W., Surget, A., Belzung, C., et al. (2021). Neuroinflammation and depression: A review. Eur J Neurosci 53, 151–171.

Troutman, T.D., Bazan, J.F., and Pasare, C. (2012a). Toll-like receptors, signaling adapters and regulation of the pro-inflammatory response by PI3K. Cell Cycle 11, 3559–3567.

Troutman, T.D., Hu, W., Fulenchek, S., Yamazaki, T., Kurosaki, T., Bazan, J.F., and Pasare, C. (2012b). Role for B-cell adapter for PI3K (BCAP) as a signaling adapter linking Toll-like receptors (TLRs) to serine/threonine kinases PI3K/Akt. Proc Natl Acad Sci U S A 109, 273–278.

Ulian-Benitez, S., Bishop, S., Foldi, I., Wentzell, J., Okenwa, C., Forero, M.G., Zhu, B., Moreira, M., Phizacklea, M., McIlroy, G., et al. (2017). Kek-6: A truncated-Trk-like receptor for Drosophila neurotrophin 2 regulates structural synaptic plasticity. PLoS Genet 13, e1006968.

Wang, C.S., Kavalali, E.T., and Monteggia, L.M. (2022). BDNF signaling in context: From synaptic regulation to psychiatric disorders. Cell 185, 62–76.

Watters, T.M., Kenny, E.F., and O’Neill, L.A. (2007). Structure, function and regulation of the Toll/IL-1 receptor adaptor proteins. Immunol Cell Biol 85, 411–419.

Wong, J.C.Y., Gokgoz, N., Alon, N., Andrulis, I.L., and Buchwald, M. (2003). Cloning and mutation analysis of ZFP276 as a candidate tumor suppressor in breast cancer. J Hum Genet 48, 668–671.

Xiang, L., Wu, Q., Sun, H., Miao, X., Lv, Z., Liu, H., Chen, L., Gu, Y., Chen, J., Zhou, S., et al. (2022). SARM1 deletion in parvalbumin neurons is associated with autism-like behaviors in mice. Cell Death Dis 13, 638.

Zhai, B., Villen, J., Beausoleil, S.A., Mintseris, J., and Gygi, S.P. (2008). Phosphoproteome analysis of Drosophila melanogaster embryos. J Proteome Res 7, 1675–1682.

Zhang, J., Tang, T., Zhang, R., Wen, L., Deng, X., Xu, X., Yang, W., Jin, F., Cao, Y., Lu, Y., et al. (2024). Maintaining Toll signaling in Drosophila brain is required to sustain autophagy for dopamine neuron survival. iScience 27, 108795.

Zhang, P., Pei, C., Wang, X., Xiang, J., Sun, B.F., Cheng, Y., Qi, X., Marchetti, M., Xu, J.W., Sun, Y.P., et al. (2017). A Balance of Yki/Sd Activator and E2F1/Sd Repressor Complexes Controls Cell Survival and Affects Organ Size. Dev Cell 43, 603–617 e605.

Zhao, J.L., Jiang, W.T., Wang, X., Cai, Z.D., Liu, Z.H., and Liu, G.R. (2020). Exercise, brain plasticity, and depression. CNS Neurosci Ther 26, 885–895.

Zheng, Y., and Pan, D. (2019). The Hippo Signaling Pathway in Development and Disease. Dev Cell 50, 264–282.

Zhou, Y., Zhou, B., Pache, L., Chang, M., Khodabakhshi, A.H., Tanaseichuk, O., Benner, C., and Chanda, S.K. (2019). Metascape provides a biologist-oriented resource for the analysis of systems-level datasets. Nat Commun 10, 1523.

Zhu, B., Pennack, J.A., McQuilton, P., Forero, M.G., Mizuguchi, K., Sutcliffe, B., Gu, C.J., Fenton, J.C., and Hidalgo, A. (2008). Drosophila neurotrophins reveal a common mechanism for nervous system formation. PLoS Biol 6, e284.

